# Catch me if you can: Capturing extracellular DNA transformation in mixed cultures via Hi-C sequencing

**DOI:** 10.1101/2022.09.16.508255

**Authors:** David Calderón-Franco, Mark C.M. van Loosdrecht, Thomas Abeel, David G. Weissbrodt

**Author notes:** Correspondence: David G. Weissbrodt, Assistant Professor, Environmental Biotechnology Section, Department of Biotechnology, Delft University of Technology, van der Maasweg 9, 2629 HZ Delft, The Netherlands.

## Abstract

Environmental microorganisms evolve constantly under various stressors using different adaptive mechanisms, including horizontal gene transfer. Microorganisms benefit from transferring genetic information that code for antibiotic resistance via mobile genetic elements (plasmids). Due to the complexity of natural microbial ecosystems, quantitative data on the transfer of genetic information in microbial communities remain unclear. Two 1-L chemostats (one control and one test) were inoculated with activated sludge, fed with synthetic wastewater, and operated for 45 days at a hydraulic retention time of 1 day to study the transformation capacity of a rolling-circle plasmid encoding GFP and kanamycin resistance genes, at increasing concentrations of kanamycin (0.01-2.5-50-100 mg L^−1^) representing environmental, wastewater, lab-selection, and gut or untreated pharmaceutical wastewater discharge environments. The plasmid DNA was spiked daily at 5 µg L^−1^ in the test chemostat. The evolution of the microbial community composition was analyzed by *16S rRNA* gene amplicon sequencing and metagenomics, and the presence of the plasmid by quantitative PCR. We used Hi-C sequencing to identify natural transformant microorganisms under steady-state conditions with low (2.5 mg L^−1^) and high (50 mg L^−1^) concentrations of kanamycin. Both chemostats selected for the same 6 predominant families of *Spirosomaceae, Comamonadaceae, Rhodocyclaceae, Rhizobiaceae, Microbacteriaceae*, and *Chitinophagaceae*, while biomass formation in the presence of kanamycin was higher with the plasmid. Hence, the antibiotic exerted the main pressure on microbial selection, while the plasmid helped these populations better resist the antibiotic treatment and grow. The kanamycin resistance gene increased in both reactors (log 7 gene copies g VSS^−1^). When higher antibiotic concentrations were applied, the GFP/16S ratio was increased, highlighting plasmids accumulation in the test reactor over time. The plasmid transformed mainly inside populations of *Bosea sp*., *Runella spp*., and *Microbacterium sp*.. This study made one significant step forward by demonstrating that microorganisms in enrichments from activated sludge biomasses can acquire exogenous synthetic plasmids by transformation.

**Graphical abstract:** 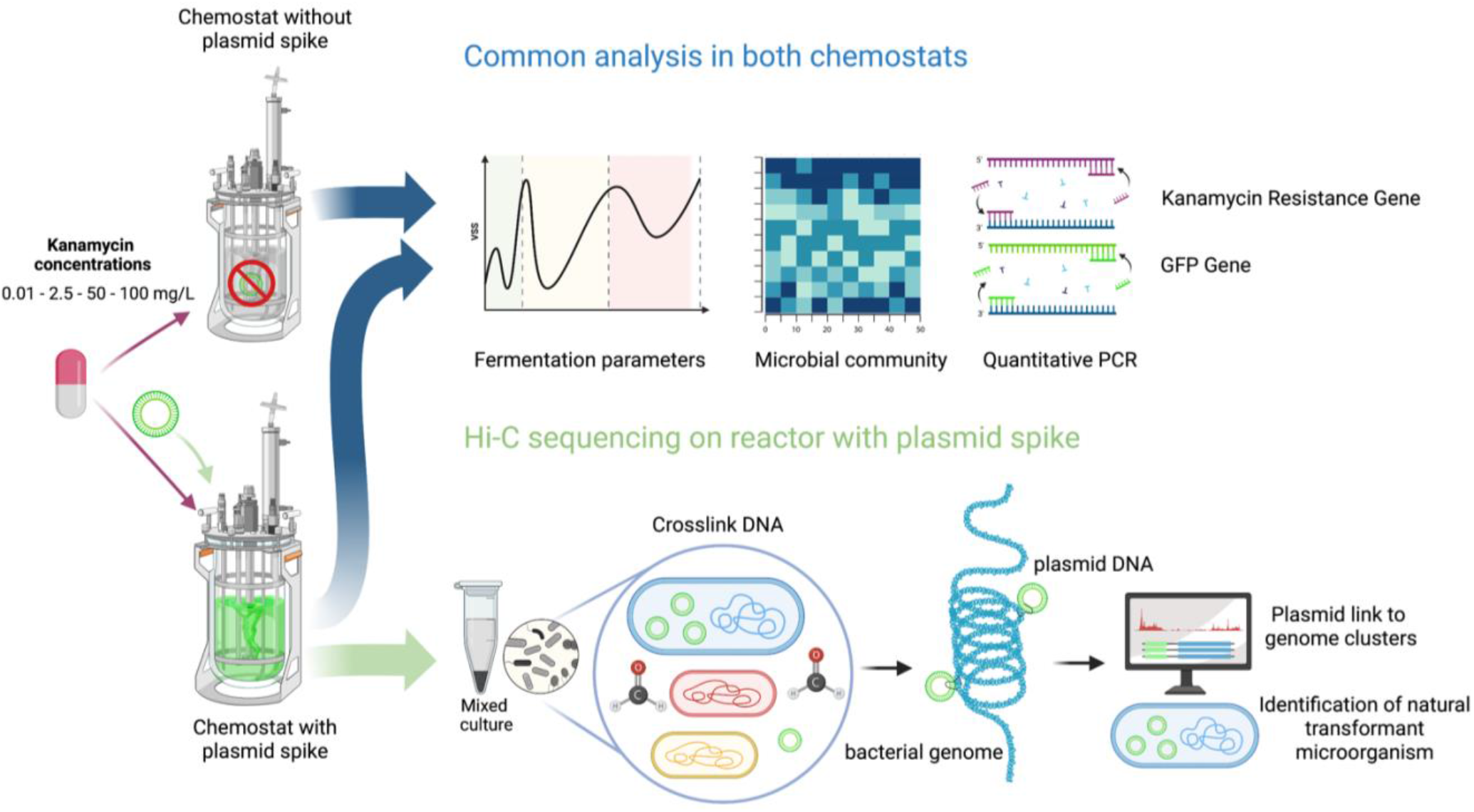

## 1. Introduction

Bacterial antimicrobial resistance (AMR) has emerged as one of the greatest public health threats of the 21^st^ century. It is estimated that by 2050, 10 million lives a year will be at risk due to the rise of drug-resistant infections by antibiotic resistance bacteria (ARB) if no mitigation efforts are engaged [1]. From the One Health context, water from places where antibiotics are highly used, such as healthcare services, agriculture, farming, households, and urban discharges, coincides collectively at wastewater treatment plants (WWTPs). This forms a broth cocktail where drugs and antimicrobial agents meet bacteria in a complex sludge.

Microorganisms in natural and man-made systems constantly evolve under environmental stressors using horizontal gene transfer (HGT) processes, such as conjugation, transduction, or transformation. Microbes benefit from transferring genetic information that code for antibiotic resistance via mobile genetic elements (MGEs) like plasmids, integrons, transposons, and conjugative and integrative elements. The quantitative elucidation of genetic information transfer by transformation in mixed microbial cultures, the environmental conditions that trigger microbial competence (seasonal changes, lack of nutrients, or antibiotic concentrations, among others), and the resistant microbial hosts, carriers, and vectors of these MGEs remain vastly unclear [2].

Recently, we highlighted that the extracellular free DNA (exDNA) in wastewater is a rich pool of MGEs (65%; 5-9 µg exDNA L^−1^) [3] that can co-localize ARGs [4]. Quantifying DNA uptake from the environment by natural transformation is of key interest [5] since it can pose a risk for the development of ARB in wastewater environments and their discharge into nature and aquatic reservoirs.

Natural transformation is the process by which bacteria can actively take up and integrate exDNA, providing a source of genetic diversity [6]. Naturally competent bacteria actively pull DNA fragments from their environment into their cells [7]. The effects of DNA uptake depend on the nutritional needs of the cells (bacteria can use exDNA as a nutrient source [8]), the presence of DNA damage, the ability of incoming DNA to recombine with chromosomal DNA, and the effects of this recombination on fitness (beneficial traits such as ARGs). Experimental demonstrations of natural competence are limited to only a few dozen species scattered across the bacterial tree and examined in pure cultures [9, 10]. Assays measuring genetic transformation are highly sensitive, but they can be done only in species that harbor a selectable genetic marker (typically an antibiotic resistance gene) and that are cultivable. Such methods fail to discover competence in complex microbial communities as present in wastewater environments, where DNA uptake rarely leads to recombination or episomal integration. The conditions that induce transformation have equally not been elucidated. Thus, one species may be mistakenly described as lacking competence since only non-competent isolates have been tested. Unraveling natural competence and transformation within microbiomes is crucial.

Most studies have so far tried to quantify gene transfer via conjugation [11–13] or transformation [14–16], either *in vitro* or *in vivo*. These approaches help quantify HGT rates under controlled conditions on bench (*in vitro*) and HGT occurrence in defined synthetic consortia using engineered or well-characterized strains cultivated as biofilms (*in vivo*). However, these methods cannot uncover direct analysis of naturally occurring HGT phenomena in microbial communities (in situ).

Despite disadvantages associated with studying HGT directly in microbial communities, such as extensive data requirements, the challenge of drawing direct cause and effect relationships between MGEs and their transfer rates [2], it is to our understanding the best approach to answer the question of who carries, transfers and can uptake ARGs in complex systems via natural transformation. Some studies have attempted to link the resistome, plasmidome, and viruses via Hi-C sequencing to the microbiome of wastewater [16] or rumen samples [17], highlighting the wide microbial diversity able to transfer genetic fragments.

Here, we studied the transformation of an exogenous synthetic rolling-circle plasmid encoding GFP and kanamycin resistance genes in an enrichment from activated sludge mixed culture fed with synthetic wastewater in chemostats exposed to increasing concentrations of kanamycin (0.01-2.5-50-100 mg L^−1^). Plasmid DNA was spiked daily in the test chemostat, while the control culture was only exposed to the antibiotic. Microbial community compositions were evaluated by *16S rRNA* gene amplicon sequencing and metagenomics, and the presence of the plasmid was analyzed by quantitative PCR. Natural transformant microorganisms were analyzed by Hi-C sequencing under steady-state conditions with 2.5 and 50 mg L^−1^ of kanamycin. We observed the natural transformation of bacteria with a synthetic plasmid coding for antibiotic resistance, in an activated sludge mixed culture.

## 2. Material and Methods

### 2.1. Mixed-culture bioreactor systems and operation

#### Chemostats

A control chemostat and a test chemostat (both of 1 L total volume and 0.7 L working volume) were operated identically in parallel, under axenic conditions (close environment), aerobically, at room temperature (23 ± 2 °C), fed with a complex synthetic wastewater (subsection 2.2.) at a hydraulic retention time (HRT) of 1 day (*i*.*e*., flowrate of 1 L d^−1^ or 0.695 mL min^−1^, and dilution rate of 1 d^−1^ or 0.0417 h^−1^), and mixed at 600 rpm by mechanical stirring. The dissolved oxygen concentration was controlled with a mass flow controller (Brooks, USA), delivering a flowrate of 0.7 L air min^−1^. Both chemostats were equipped with oxygen sensors (AppliSens, Poland), thermometers, and pH probes (Mettler Toledo, USA). pH was maintained at 7.0 ± 0.5 by addition of HCl or NaOH at 1 mol L^−1^ each. All samples were taken in sterility using a tube welder (Tekyard, USA). The off-gas from the reactor was filtered-sterilized before release. The effluent was collected in a closed vessel and always autoclaved before discarding. A schematic representation of the equipment is provided in **Figure 1**.

**Figure 1.**
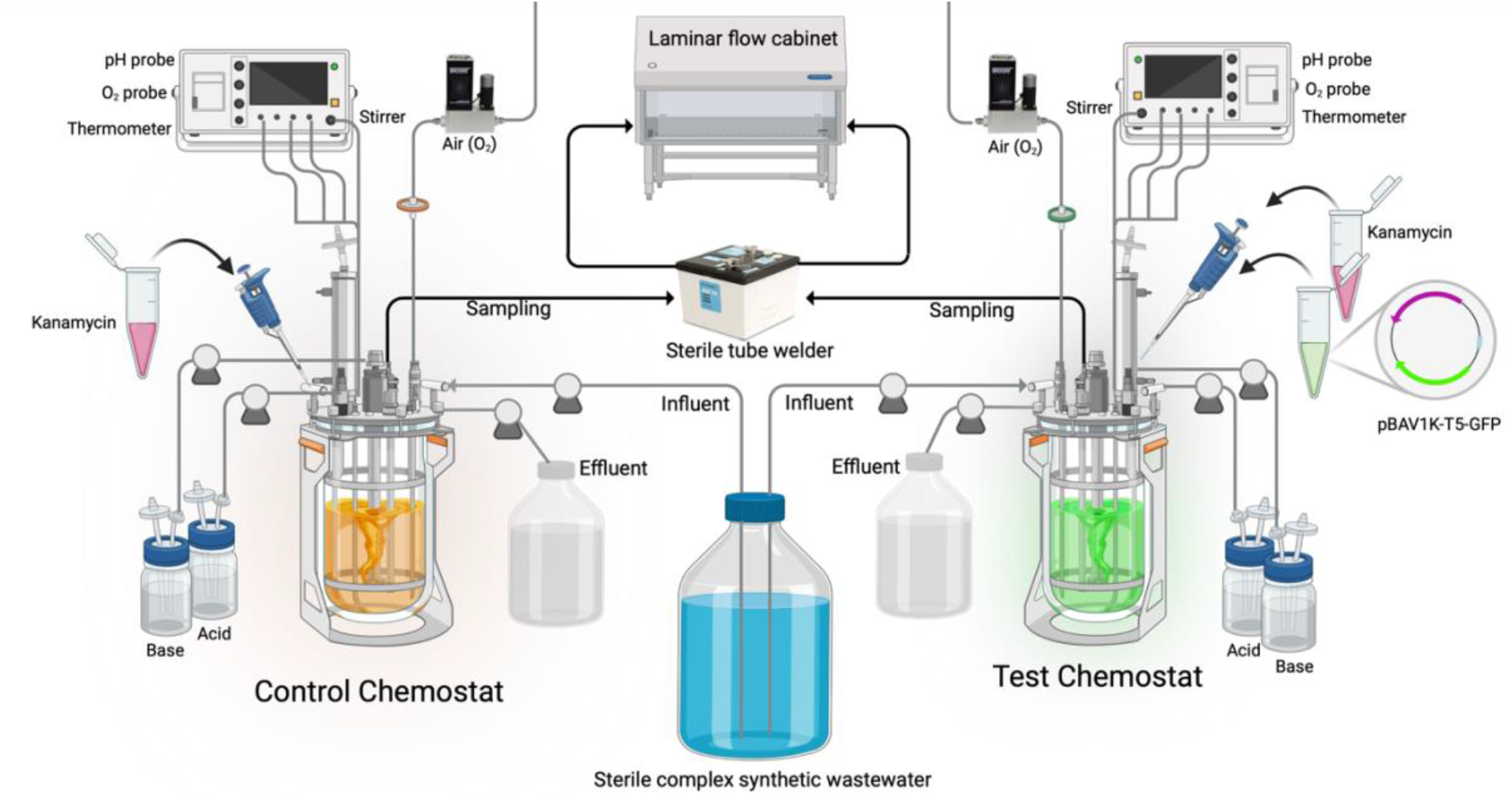
Schematic representation of the set containing two continuously stirred bioreactors (chemostats): the control chemostat, where only kanamycin was spiked, and the test chemostat, where kanamycin and the test plasmid (pBAV1K-T5-GFP) were daily spiked. This experimental setting was embedded in a biosafety level II laboratory, where samples had to be extracted via a sterile tube welder and handled under sterility in a laminar flow cabinet.

#### Inoculum

The reactors were inoculated at a low initial concentration of biomass of 0.043 g VSS L^−1^, by seeding 10 mL of activated sludge (3 g VSS L^−1^) collected from WWTP Amsterdam-West (The Netherlands), which is operated for complete biological nutrient removal. The inoculated biomass was equilibrated to the chemostat operation in both reactors over 5 HRTs in the absence of antibiotics.

#### Antibiotic supply

After acclimation, the reactors were run for 46 days under increasing concentrations of the model antibiotic kanamycin (Thermo Fisher Scientific, USA) from 0.01 to 2.5, 50, and 100 mg L^−1^. Antibiotic loadings were changed after maintaining each condition over 8-10 HRTs. Kanamycin was spiked daily in both chemostats simultaneously (**Figure S1**).

#### Plasmid spikes

In the test reactor only, the rolling circle plasmid pBAV1K-T5-GFP containing genes coding for resistance to kanamycin and a green-fluorescent protein (GFP) as a reporter gene [17] was spiked daily (5 µg L^−1^) for 37 days. The Monarch Plasmid Miniprep Kit (New England BioLabs, USA) was used to isolate plasmid DNA. This plasmid backbone has already been used for natural transformation in inter-species studies [14]. The plasmid was not spiked over the additional last 10 days of the experiment, to assess if the plasmid would remain accumulated in the mixed culture or washed out due to the continuous operation. The spikes of antibiotics and plasmid were injected through sterile luer connections.

### 2.2. Complex synthetic wastewater composition

A complex synthetic wastewater was prepared according to [18] and previously applied for conjugative experiments [11], to obtain a controlled composition mimicking real wastewater. The detailed influent composition is shown in **Table S1** and **Table S2** in the **Supplementary Material**.

Briefly, this medium was composed of 1/3 volatile fatty acids (1/6 acetate + 1/6 propionate), 1/3 soluble and fermentable substrates (1/6 glucose, 1/6 amino acids), and 2 particulate substrates (1/6 peptone, 1/6 starch) in equal equivalents of chemical oxygen demand (COD).

Amino acids were composed of L-alanine, L-arginine, L-aspartic acid, L-glutamic acid, L-leucine, L-proline, and glycine in COD equivalents. Particulate substrates were peptone from casein, digested with trypsin (Carl Roth, Germany), and starch made from wheat (Merck Sigma, Germany).

Nitrogen was supplied as a combination of soluble ammonium chloride and nitrogen from the aforementioned amino acids and peptone. Phosphorus was composed of soluble orthophosphate.

### 2.4. Analytical methods for the measurements of substrates and biomass concentrations Chemical analyses of the liquid phase

The mixed liquor of the chemostats was sampled as volumes of 11 mL from the effluent in triplicates at a sample frequency every 2 days. 1 mL was centrifuged at 6000 x g 1 min and supernatants filtered through 0.2 µm PVDF membrane filters (Pall, USA), and the filtrates were used for chemical analyses of the liquid phase. Acetate, propionate, and glucose concentrations were measured using a high-performance liquid chromatograph (HPLC; Vanquish™ System, Thermo Fisher Scientific, USA) using an Aminex HPX-87H column (BioRad, USA) maintained at 59°C and coupled to an ultraviolet detector at 210 nm (Waters, USA) and a refractive index detector (Waters, USA). A solution of phosphoric acid at 1.5 mmol L^−1^ was used as eluent. Total nitrogen (5-40 mg L^−1^ TN range) and phosphate (0.5-5.0 mg L^−1^ PO ^3-^-P) were measured with colorimetric-spectrophotometric cuvette tests (Hach-Lange, USA).

#### Biomass analyses

The concentrations of total suspended solids (TSS) and volatile suspended solids (VSS) of the mixed liquors were analyzed according to Standard Methods [19]. The other 10 mL obtained from the mixed liquor every two days for chemical analyses were used for TSS and VSS measurements.

### 2.5. Preliminary control of plasmid transformation in pure cultures

*Escherichia coli* K12 and *Bacillus subtilis* str. 168 were used as preliminary controls to verify that the plasmid could transform and express in Gram-negative and Gram-positive microorganisms. *E. coli* cells were electroporated, and *B. subtilis* was transformed with a starvation-induced method [20] with the plasmid and plated in Luria-Bertani (LB) medium with kanamycin (50 mg L^−1^) to select for positive transformants. Details on electroporation and starvation-induced transformation can be found in **Supplementary Material**. Microscopic bright field and fluorescent pictures of positively transformed *E. coli* and *B. subtilis* can be found in **Figure S2**.

### 2.6. Quantitative PCR analysis of kanamycin resistance and GFP marker genes

All qPCR reactions were conducted in 20 µL, including IQ™ SYBR green supermix BioRad 1x. The sets of forward and reverse primers used to amplify the green fluorescent protein (GFP) gene and the kanamycin resistance (*kanR*) gene were retrieved from [17] and summarized in **Tables 1 and S3**. A volume of 2 µL of DNA template was added to each reaction, and the reaction volume was completed to 20 µL with DNase/RNase free Water (Sigma Aldrich, UK). All reactions were performed in a qTOWER3 Real-time PCR machine (Westburg, DE) according to the following PCR cycles: 95°C for 5 min followed by 40 cycles at 95°C for 15 s and 60°C for 30 s.

**Table 1.**
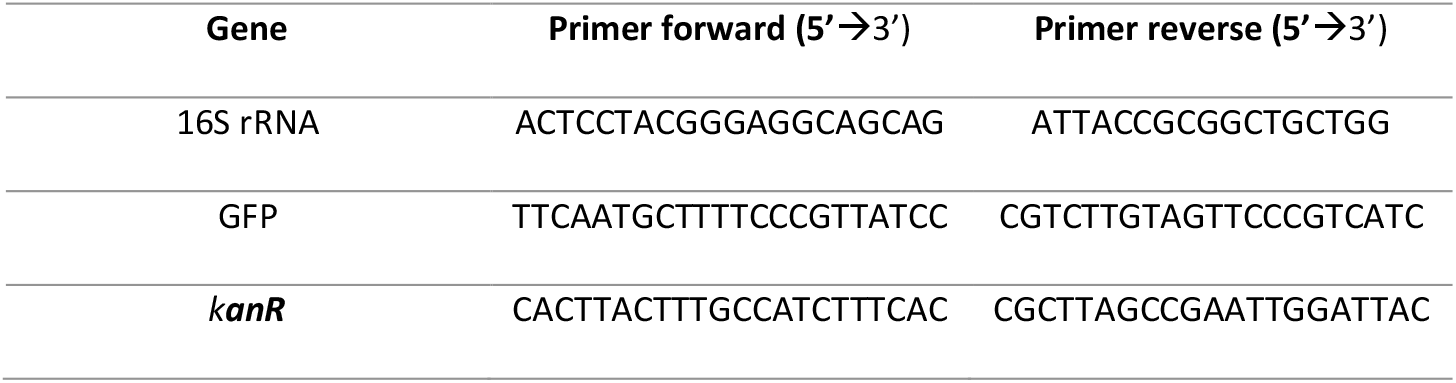
Primers used for qPCR analyses

To check the specificity of the reaction, a melting curve was performed from 65 to 95°C at a temperature gradient of +0.5°C (5 s)^−1^. Synthetic DNA fragments (IDT, USA) containing each target gene were used as a positive control to create the standard curves. Serial dilutions of gene fragments were performed in sheared salmon sperm DNA at 5 µg mL^−1^ (m/v) (Thermofisher, LT) diluted in Tris-EDTA (TE) buffer at pH 8.0. Every sample was analyzed in technical triplicates. Standard curves were included in each PCR plate with at least six serial dilution points and technical duplicates. An average standard curve based on a standard curve from every run was created for every gene set. Gene concentration values were then calculated from the standard curve mentioned above.

### 2.7. Microbial populations dynamics by 16SrRNA gene amplicon sequencing

Changes in compositions of the microbial communities of the two chemostats during antibiotic regime shifts were analyzed by amplicon sequencing. Volumes of 5 mL of mixed liquors were taken at the end of each antibiotic concentration period (*i*.*e*., at steady states) in 15 mL Falcon tubes. Cell pellets were obtained by centrifugation at 6000 x g during 1min. DNA was extracted from the samples’ cell pellets using the PowerSoil microbial extraction kit (Qiagen Inc., Germany), following manufacturer’s instructions. The DNA content of the extracts was quantified using a Qubit 4 (Thermo Fisher Scientific, United States). The DNA extracts were preserved at −20°C pending amplicon sequencing analyses.

The DNA extracts were sent to Novogene Ldt (Novogene, Hong Kong) for the V3-V4 16SrRNA gene hypervariable regions (position 341-806) on a MiSeq desktop sequencing platform (Illumina, San Diego, USA). The raw sequencing reads were processed by Novogene Ltd and quality filtered using the QIIME software [21]. Chimeric sequences were removed using UCHIME [22], and sequences with ≥97% identity were assigned to the same operational taxonomic units (OTUs) using UPARSE [23]. Each OTU was taxonomically annotated using the Mothur software against the SSU rRNA database of the SILVA Database [24]. The heatmap of relative abundances was generated using the R package “ampvis2” v2.7.31 [25].

### 2.8. Microbiome profiling by metagenomics

The same biomass samples selected for Hi-C metagenomics sequencing (collected at 2.5 and 50 mg Kan L^−1^; see §2.9 below) were sequenced in parallel by conventional metagenomics to profile their microbiomes at high resolution. Samples were submitted to and sequenced by Phase Genomics (USA). Both the shotgun library and the Hi-C library were sequenced on the same platform. The details are given in section §2.9.

Classification with Kraken2.0 [26] was performed on pair-end mode on the quality-controlled short reads, using the Microbial Database for Activated Sludge (MiDAS) [27]. The taxonomic classification outcomes from Kraken2.0 were converted into a BIOM file using the kraken-biom [28] tool to explore metagenomics classification datasets via the “MicrobiotaProcess” package v1.6.6. in R [29].

### 2.9. Hi-C sequencing

Hi-C libraries were prepared as explained in [30] and summarized in **Figure 2**.

**Figure 2.**
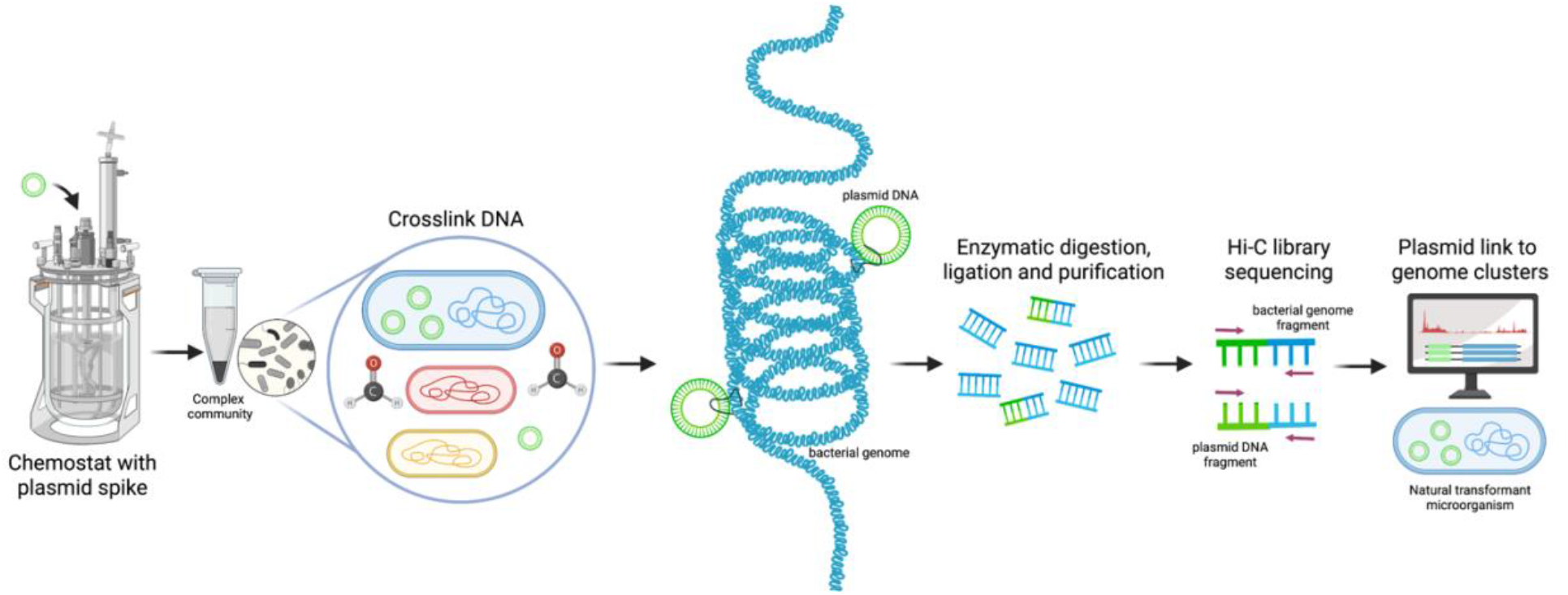
Schematic representation of the Hi-C deconvolution process in which the microbial community at a particular time point is crosslinked with formaldehyde before cell lysis, linking plasmids to bacterial genomes. DNA extract is digested enzymatically, biotinylated, ligated, and purified. To generate the Hi-C library, the resultant fraction is sequenced and used to create Hi-C links used to deconvolute contigs into genome clusters, including chromosomes and plasmids.

#### Biomass sampling

Biomass samples were collected as volumes of 10 mL of mixed liquor at the end of the antibiotic treatment periods at 2.5, and 50 mg Kan L^−1^ from the test reactor spiked with the plasmid and transferred on ice. The biomass was aliquoted in two 1.5 mL Eppendorf. One sample was used for Hi-C library generation, while the other was used for standard metagenomic library preparation.

#### Biomass sample conditioning for Hi-C analysis

A volume of 1.5 mL of the collected biomass sample was resuspended using 13.5 mL of solution of a commercial 1% formaldehyde-phosphate-buffered saline (F-PBS) (Alfa Aesar - Thermo Fischer Scientific, USA) in a 15 mL Falcon tube under biosafety level II conditions. Formaldehyde was used to generate covalent links between spatially adjacent genetic segments (**Figure 2**). The resuspended sample was incubated at room temperature for 30 min with periodic mixing (every 5 min) before adding glycine at a final concentration of 1 g/100 mL to quench the reaction and further incubating the mixture at room temperature for 15 min with periodic mixing. The last step involved a series of three spin down (6000 x g, 1 min) and rinses with PBS of the pellet. Briefly, the sample was spun down for 2 min at 6000 x g, rinsed with PBS, and spun down again (5 min at 6000 x g) before removing the supernatant. The final pellet was kept frozen at −20 °C.

#### Hi-C library preparation and sequencing

Hi-C libraries were prepared with the Phase Genomics ProxiMeta Hi-C v4.0 Kit using the manufacturer-provided protocol [31]. Briefly, the intracellular DNA pool (comprising the genomic DNA and the accessory genome, i.e., MGEs) of the microbial cells present in each of the two selected samples were crosslinked using the formaldehyde solution. These pre-treated biomass samples were submitted to Phase Genomics for library preparation and sequencing. Cells were bead-beated to release the cross-linked DNA. The released cross-linked DNA was digested using the Sau3AI and MlucI restriction enzymes simultaneously, and proximity ligated with biotinylated nucleotides to create chimeric molecules composed of fragments from different regions of genomes and plasmids that were physically proximal *in vivo*. Proximity ligated DNA molecules were pulled down with streptavidin beads and processed into an Illumina-compatible sequencing library. Separately, using an aliquot of the original samples, DNA was extracted with a ZYMObiomics DNA miniprep kit (Zymo Research, USA) and a metagenomic shotgun library was prepared using ProxiMeta library preparation reagents.

Sequencing was performed on an Illumina NovaSeq generating PE150 read pairs for both the shotgun libraries (§2.8) and the Hi-C libraries (§2.9) obtained from the aliquots of each of the two biomasses collected at 2.5 and 50 mg Kan L^−1^. Hi-C and shotgun metagenomic sequencing files were uploaded to the Phase Genomics cloud-based bioinformatics platform for subsequent analysis.

### 2.10 Processing of shotgun metagenomics and Hi-C metagenomics datasets

#### Quality control of sequenced reads

After sequencing, datasets containing 4 paired-end read samples with an average of 78M reads for the Hi-C samples, and 160M reads for the non-crosslinked samples (i.e., shotgun) were obtained. The quality of the Illumina reads was assessed using FastQC version 0.11.9 with default parameters [32]. Shotgun reads were filtered and trimmed for quality and normalized using fastp v0.19.6 [33].

#### Assembly of shotgun sequence reads

The trimmed reads were assembled into contigs using MEGAHIT v1.2.9 [35] for the resistome analysis. The trimmed reads were assembled into contigs using metaSPAdes version 3.14.1 [34] on meta mode on default parameters for following discordant reads analysis (to quantify interactions plasmid:bacteria). We used these two assemblers to verify which one resulted in positive results for plasmid-host detection (only metaSPAdes displayed positive results).

#### Processing of the Hi-C reads

Each set of Hi-C reads was mapped to the metagenomic assemblies to generate a SAM file containing the information of the assembly and the Hi-C links. Mapping was done using the Burrows-Wheeler alignment tool BWA-MEM v0.7.17-r1188 [36]. During mapping with BWA MEM, read pairing and mate-pair rescue functions were disabled and primary alignments were forced to be aligned with the lowest read coordinate (5’ end) (options: -5SP) [37]. SAMBLASTER v0.1.26 [38] was used to flag PCR duplicates, which were later excluded from the analysis. Alignments were then filtered with samtools v1.13 [39] using the -F 2304 filtering flag to remove non-primary and secondary alignments.

#### Deconvolution of the Hi-C data for aminoglycoside resistome analysis

Metagenome deconvolution was performed with ProxiMeta [40, 41], creating putative genome and genome fragment clusters (**Figure S14-S15**). Clusters (also known as metagenome-assembled genomes or MAGs) were assessed for quality using CheckM v1.1.10 [42] (>90% completeness, <10% contamination) and assigned preliminary taxonomic classifications with Mash v2.3 [43]. NCBI plasmid database was used to identify which contigs had plasmid or genomic DNA as the origin, and NCBI AMRFinderPlus software (v3.10.5) [44] was used to annotate aminoglycoside resistance genes using the NCBI AMRFinder database. AMR genes and plasmids were annotated on all contigs in the assembly. Hi-C signal was used to associated plasmids with their hosts. Thus, if a plasmid was associated with a host and had an AMR gene, the event was annotated as an AMR gene conveyed to the host via a plasmid. If an AMR gene was found on a binned contig that was not annotated as a plasmid, it was classified as an AMR gene originating from genomic DNA [45]. The taxonomic trees were generated with ‘‘ggtree” package v3.2.1 in R [46].

#### Plasmid transformation events detection from Hi-C data using discordant-reads analysis

Both ProxiMeta platform and bin3C v0.1.1. [37] tool were used for the generation of clusters to look for pBAV1K-T5-gfp integration. These available methods were not sensitive enough for detecting Hi-C links by identifying plasmid-contig (host) events. We therefore implemented a discordant reads analysis to quantify the interactions obtained from crosslinking the DNA pool between two genetic sequences that are not necessarily consecutive in the bacterial genome and accessory genome, within cells of the mixed cultures. A detailed explanation of the discordant read analysis, workflow, and visualization of transformation events can be found in **Supplementary Material**.

## 3. Results

### 3.1. Plasmid spike had a beneficial effect on biomass growth under acute antibiotic treatment

The biomass in both chemostats grew at a growth rate (µ) of 0.042 h^−1^ (*i*.*e*., equal to the dilution rate applied) under environmental antibiotic concentrations (0.01-2.5 mg Kan L^−1^) **(Figure 3)**. When higher antibiotic pressures were applied (50 – 100 mg Kan L^−1^), a significantly higher biomass formed (max Δ150 mg VSS L^−1^) when the plasmid was spiked in the broth. Uptake of plasmid DNA and the kanamycin resistance gene expression can confer a better resistance to higher antibiotic concentrations. Interestingly, the control reactor treated with a high concentration of kanamycin of 50 mg L^−1^ displayed an abrupt decrease in culture viability compared to the test reactor amended both with the antibiotic and the plasmid. The highest antibiotic concentration (100 mg L^−1^) was eventually selected for biomass able to grow in the control reactor. Details on operation performances are given in **Figure S3**.

**Figure 3.**
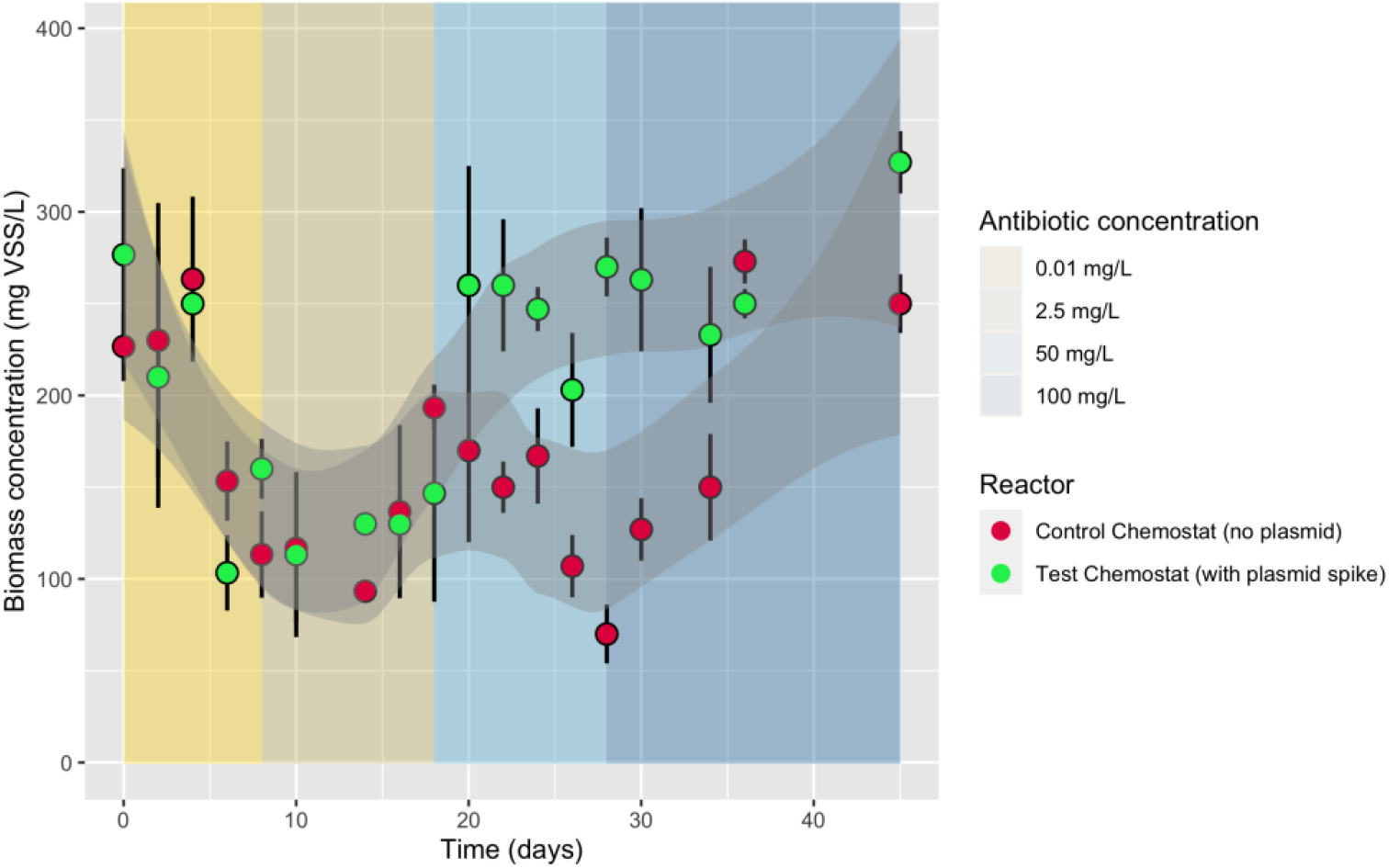
Daily-averaged values for volatile suspended solids (VSS) from both reactor control and reactor with free-floating plasmid over the whole operation time the experiment was conducted (45 days). Kanamycin concentrations are displayed as background-colored sections: 0.01 mg L^−1^, 2.5 mg L^−1^, 50 mg L^−1^, 100 mg L^−1^.

### 3.2. Microbial community compositions under increasing antibiotic pressure

Microbial population composition **(Figure 4a)** was analyzed by 16S rRNA gene amplicon sequencing from both the control (RC) and test (RT) reactors at the end of each period of increasing kanamycin loading. After inoculation, the reactors were acclimatized for 10 days to chemostat operation prior to antibiotic supply and plasmid spikes (day 0 is the moment of the start of antibiotic addition). The same predominant bacterial populations were enriched in both reactors. Rather than the plasmid spikes, the antibiotic supply exerted the strongest selection pressures on the microbial communities. There was a decrease in diversity with increased antibiotic dosage as it enriched for 5-6 families of *Spirosomaceae, Comamonadaceae, Rhodocyclaceae, Rhizobiaceae, Microbacteriaceae*, and *Chitinophagaceae*.

**Figure 4.**
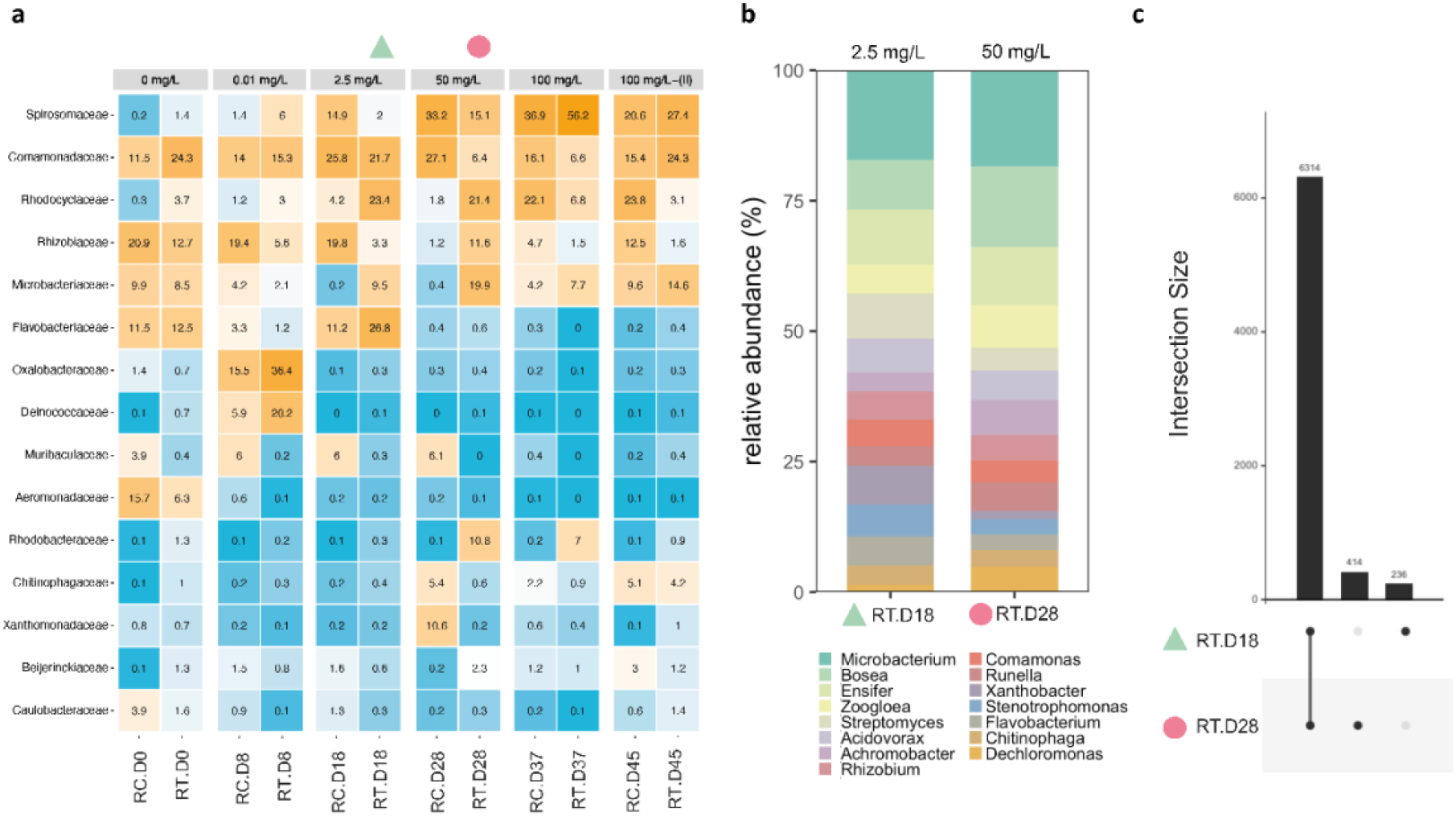
**(a)** Heatmaps from 16SrRNA amplicon sequencing showing bacterial family compositions across the operation in both chemostats grouped per kanamycin concentration (0 mg L^−1^, 0.01 mg L^−1^, 2.5 mg L^−1^, 50 mg L^−1^, 100 mg L^−1^ with spiked plasmid and 100 mg L^−1^ –(II) without spiked plasmid). The different color intensities represent the relative bacterial family abundance in each population. **Note**: **RC** = Reactor Control without spiked plasmid. **RT** = Reactor Test with spiked plasmid. (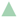: Day 18 – 2.5 mg L^−1^) and 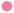: Day 28 – 50 mg L^−1^) represent the samples from the reactor test that were also sent to analyze for Hi-C sequencing. **(b)** Microbiome analysis from the samples sent for Hi-C sequencing at the genus level. **(c)** UpSet plot showing microbiome intersections between day 18 and day 28 from the test reactor.

The ratios of GFP and *kanR* over 16S rRNA strongly increased in the test system under high antibiotic concentration. Therefore, biomass samples were collected from the test reactor during the stationary periods at 2.5 (day 18; Test Reactor Day 18, i.e., sample RT.D18) and 50 (day 28; RT.D28) mg L^−1^ of kanamycin and sequenced for metagenomics and Hi-C analyses to uncover the resistome, mobilome, and transformation events in the mixed culture. The metagenome of these two samples RT.D18 (2.5 mg Kan L^−1^) and RT.D28 (50 mg Kan L^−1^), shared 90.6% of reads that mainly affiliated with *Microbacterium, Bosea, Ensifer, Zoogloea*, and *Streptomyces* genera (**Figure 4b-c**).

The list of MAGs recovered by aligning Hi-C reads against the shotgun assembly, and their quality, both RT.D18 and RT.D28, can be found in **Figure S14** and **Figure S15**, respectively.

### 3.3. The spiked plasmid accumulated in the system at high antibiotic concentrations

The kanamycin resistance gene was quantified by qPCR during the whole operation of both the control and test reactors (**Figure 5**). A kanamycin-resistant enrichment culture developed with the increasing antibiotic dosage in both reactors. The kanamycin gene was obtained by the microorganisms either via natural transformation from the spiked plasmid pBAV1K-T5-GFP or endogenously present in the activated sludge inoculum.

**Figure 5.**
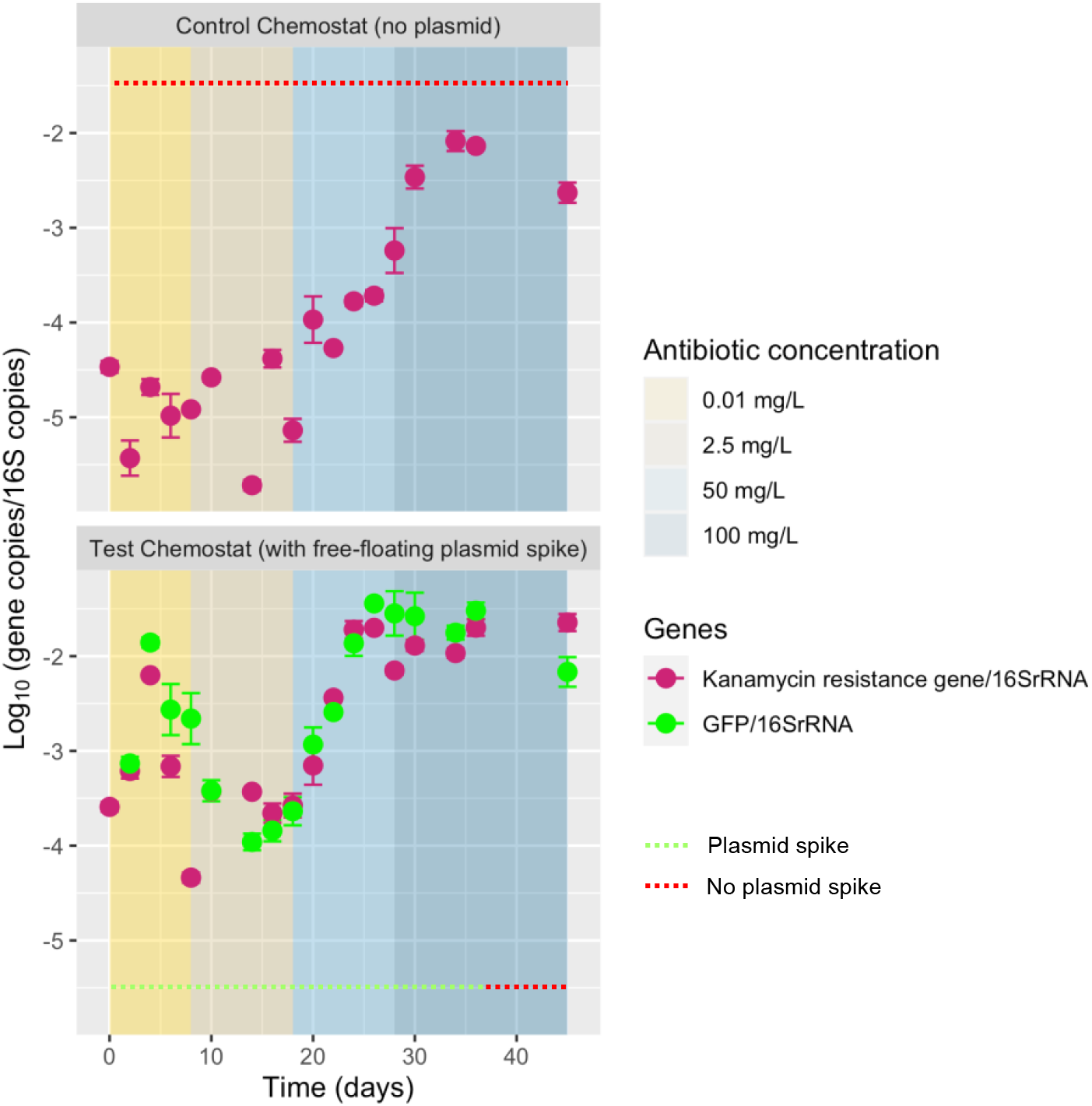
Relative abundance of green fluorescent protein (GFP) gene and Kanamycin resistance gene from plasmid pBAV1K-T5-GFP relative to 16S rRNA gene in every sampling point. Only data from the reactor with free-floating plasmid displayed as non-detected values were detected for reactor control. Kanamycin concentrations are displayed on colored-background sections: 0.01 mg L^−1^, 2.5 mg L^−1^, 50 mg L^−1^, 100 mg L^−1^. The plasmid was not spiked anymore in the test reactor (after day 36).

The spiked plasmid integrated the kanamycin resistance gene and a GFP gene. qPCR was also performed on the GFP gene. The GFP gene was detected in the test reactor spiked with the plasmid but not in the control reactor (**Figure 5, upper panel**). Hence, the plasmid was exclusively present in the test reactor. Reporter GFP fluorescence measurements could not be used as validation due to the sample complexity or lack of expression.

Higher antibiotic concentrations resulted in an enrichment of the ratio of GFP to 16S rRNA genes (GFP/16S ratio) present in the test mixed culture from −3.5 to −1.5 log gene copies (GFP and *kanR*, respectively)/16S rRNA **(Figure 5)**. Plasmid accumulation occurred either by uptake by bacteria and replication during growth or adsorbed to extracellular polymeric substances (EPS)[47]. After 18 days of exposure to the highest kanamycin concentration of 100 mg L^−1^, no plasmid was spiked in the test culture for additional 8 days. The resulting GFP/16S ratio did not decrease substantially, supporting the hypothesis that the plasmid had been integrated into the mixed culture’s microbial population.

### 3.4. Increasing kanamycin concentrations promoted plasmid mobility and aminoglycoside resistance genes Hi-C interactions

The impact of increasing concentrations of kanamycin (aminoglycoside antibiotic family) was not only investigated for the abundance of the kanamycin resistance gene spiked with the synthetic plasmid, but also for the broader presence of aminoglycoside resistance genes of genomic or natural plasmid origins, and their association in bacterial hosts (**Figure 6)**.

**Figure 6.**
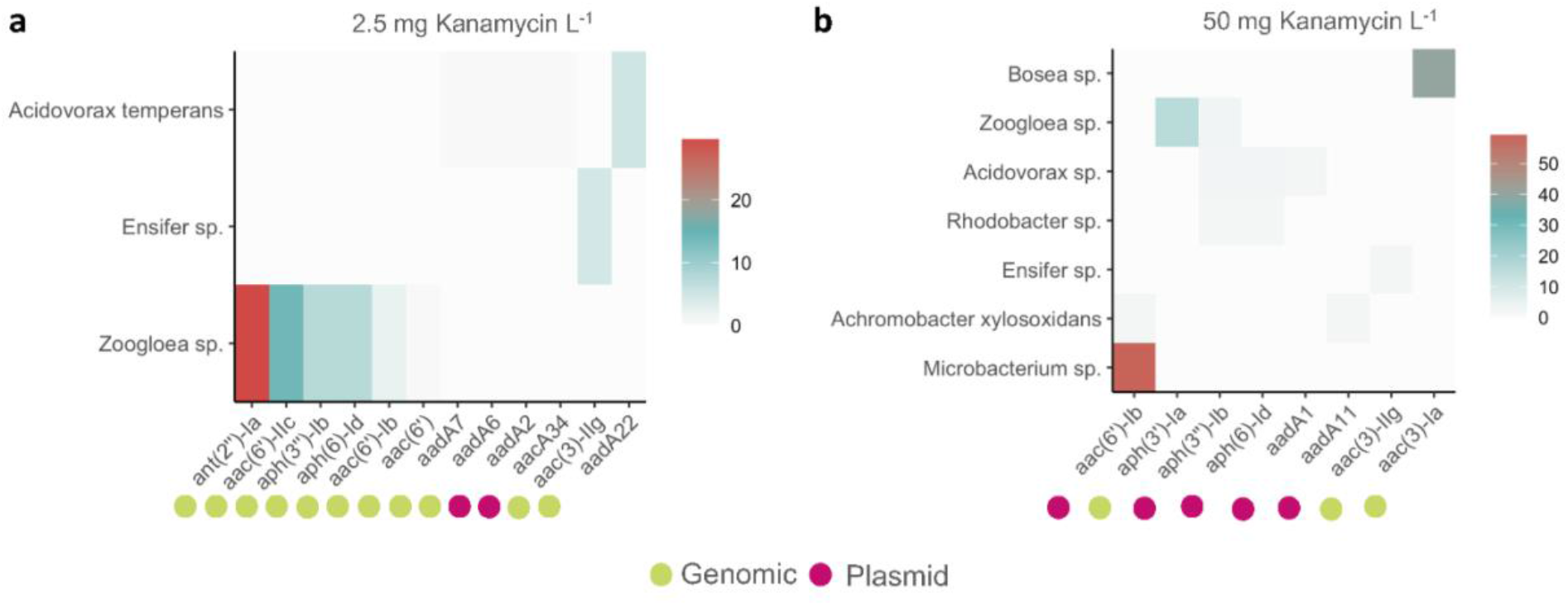
The number of Hi-C interactions between aminoglycoside resistance genes and bacterial hosts when **(a)** 2.5 mg Kanamycin L^−1^ and **(b)** 50 mg Kanamycin L^−1^ were applied to the test chemostat. Contigs containing aminoglycoside resistance genes were classified to determine their origin: genomic or plasmid DNA. Colors represent the number of Hi-C interactions normalized by the contig length (kb). Neighbor-joining trees of the different aminoglycoside resistance genes and their alignment can be found in **Figure S4-S5**.

The higher kanamycin concentration (50 *vs*. 2.5 mg L^−1^) doubled the number of bacterial hosts genomic sequences’ interacting with aminoglycoside resistance genes (3 vs 7 bacterial hosts). The maximum number of Hi-C interactions between genomic sequences and aminoglycoside resistance genes (expressed as *gene:bacterial host*) in the mixed culture also doubled: *ant(2’’)-Ia*:*Zoogloea* sp. (29 Hi-C interactions kb^−1^; **Figure 6a**) vs. *aac(6’)-Ib*:*Microbacterium* sp. (59 Hi-C interactions kb^−1^; **Figure 6b**). Thus, more bacterial hosts contained genes coding for aminoglycoside hydrolyzing enzymes and more copies of these genes (that can occur both on the genomic DNA and on naturally occurring plasmids) were detected under high antibiotic pressure.

There was a shift in the origin of the resistance genes: genomic or plasmid DNA. The low kanamycin concentration (2.5 mg L^−1^) selected mainly for *Zoogloea* sp. which harbored multiple aminoglycoside resistance genes integrated into its genome. Higher concentrations of kanamycin promoted mobility of aminoglycoside resistance genes encoded in plasmids. We detected multiple aminoglycoside resistance genes encoded in plasmids. Some of genes detected in plasmids were found in multiple bacteria such as the aminoglycoside 3’-phosphotransferase gene (*aph(3’’)-Ib)* in *Zoogloea* sp., *Acidovorax* sp., and *Rhodobacter* sp. **(Figure 6b)**.

### 3.5. Discordant reads analysis allowed the quantification of spiked plasmid transformation in bacterial species

Hi-C sequencing results show how difficult it is to capture a transformation event when a specific plasmid is spiked in a mixed culture. Bioinformatic tools available for the analysis of Hi-C libraries, such as bin3c [37] or commercial ProxiMeta platform [48], were not suited to identify transformation events of one specific plasmid considering the complex diversity of plasmids and other MGEs present in the samples. Therefore, we performed a discordant reads analysis of the aligned Hi-C reads against the shotgun metagenome assembly. The discordant reads analysis quantifies the interactions obtained from crosslinking the DNA pool between two genetic sequences that are not necessarily consecutive in the genome of a bacterial population present in the mixed culture. Synthetic constructs consisting of the plasmid sequence, the non-coding spacer DNA, and the metagenome contigs with higher Hi-C links with plasmid contigs were generated to verify the transformation of the spiked plasmid inside bacteria and its integration into their genome or its episomal presence in their cytoplasm **(Figure 7a, Figure S7)**. Hi-C metagenomes were sequenced at antibiotic pressures of 2.5 mg L^−1^ (sample RT.D18) and 50 mg L^−1^ (RT.D28) of kanamycin. No plasmid sequence was retrieved from the raw reads and the assembly of sample RT.D18. At this time point, the plasmid was either present at a very low concentration (which corresponds to the lowest plasmid/16S ratio detected by qPCR; **Figure 4**), but metagenomics was not sensitive enough to detect it.

**Figure 7.**
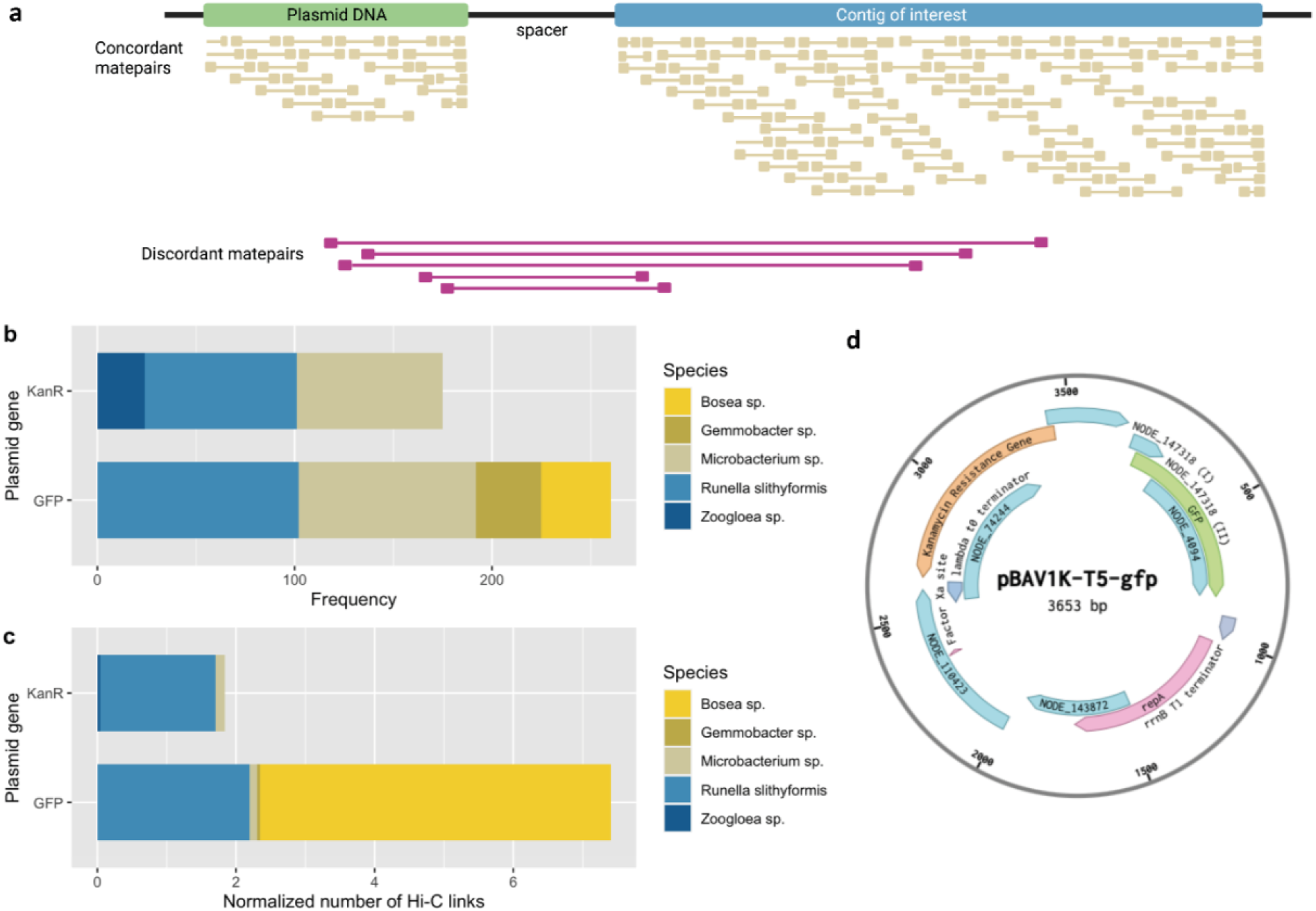
**(a)** Schematic representation of the discordant read analysis performed on contigs interacting with plasmid contigs, where synthetic constructs were generated to quantify the number of interactions from the Hi-C library alignment. **(b-c)** Taxonomic assignment of contigs that were linked to contigs harboring the information of the spiked pBAV1K-T5-GFP plasmid by frequency and by a normalized number of Hi-C links (#events divided by the length of the interacting contig). **(d)** Plasmid map containing the contigs used for the discordant read analysis. The ***kanR*** information in **a-b** is a combination of the number of interactions between the Kanamycin Resistance Gene with the NODE_74244 and NODE_147318 (I). Likewise, the GFP information is a combination of the number of interactions between the **GFP** gene with the NODE_147318 (II) and NODE_4094.

Conversely, the assembly RT.D28 remarkably harbored multiple contigs containing genetic information from the spiked plasmid. This highlights that synthetic plasmids can end up inside microbial populations of an activated sludge mixed culture through transformation. The genomic DNAs of the Gram-negative *Bosea* sp., *Runella* spp., *Gemmobacter* sp., *Zoogloea* sp., and of the Gram-positive *Microbacterium* sp. were cross-linked with genetic fragments coming from the spiked plasmid (**Figure 7b**). *Runella slythiformis* contigs displayed the highest frequency of Hi-C links (102), followed by *Microbacterium oxydans* (90) (**Figure 7b**). *Bosea* sp. had the highest number of Hi-C links per kb of contig sequence length (5.1 Hi-C links kb^−1^), followed by *Runella* spp. (2.2 Hi-C links kb^−1^) (**Figure 7c**). Genus associated to the previous bacteria had a relative fold increase of 2-4 times when compared the microbiome from 2.5 and 50 mg Kan L^−1^. *Microbacterium* increased from 5.4 mg VSS L^−1^ to 14.6 mg VSS L^−1^; *Bosea* increased from 3 mg VSS L^−1^ to 12.4 mg VSS L^−1^; *Runella* increased from 1.6 mg L^−1^ to 3.3 mg L^−1^; *Zoogloea* increased from 1.74 mg VSS L^−1^ to 6.6 mg VSS L^−1^.

Only contigs interacting with the kanamycin resistance gene (*kanR*) or the GFP gene were used to quantify Hi-C links (**Figure 7d**). This also informs that microorganisms from different classes, such as *Alphaproteobacteria, Actinobacteria, Bacteroidetes*, and *Betaproteobacteria*, were transformed in the mixed culture with this rolling-circle plasmid (**Figure S8**). Visualization examples of cross-linking events with full plasmid are given in **Figures S9-S12**. They strongly confirmed that *Bosea sp*. and *Runella* spp. took up the plasmid from the environment (**Figure S8 and Figure S9**). Interactions visualization was less clear for the other microbial populations with detected Hi-C links.

These results show that (synthetic) plasmids can transform into different microorganisms of multiple classes in activated sludge mixed cultures when enough selection pressures (high antibiotic concentrations) are introduced into the system.

## 4. Discussion

### 4.1. The growth of the activated sludge biomass under high antibiotic concentrations was promoted by plasmid accumulation

The microbial enrichment culture was able to adapt and substantially grow under high antibiotic concentrations (≥50 mg Kan L^−1^) concomitantly to plasmid accumulation in the test reactor (**Figures 3-4**). At high antibiotic loading, the biomass grew with higher yields in the test reactor than in the control reactor, which was not spiked with the plasmid.

Microorganisms transformed and selected at high concentrations of kanamycin could improve their fitness and resist antibiotic treatment. The plasmid kanamycin resistance gene confers resistance against the antibiotic, allowing bacterial cells to withstand and grow under high kanamycin concentrations. The kanamycin resistance gene was detected in the test reactor spiked with the plasmid designed with kanR and in the control reactor that did not receive the plasmid. The qPCR results showed that microorganisms with kanamycin resistance genes were already in the activated sludge inoculum collected at the full-scale WWTP. Genes coding for resistances to aminoglycosides (such as kanamycin) have been identified in the influents, sludges, and effluents of WWTPs [49–51].

The most abundant aminoglycoside resistance genes in the system were *aac(6’)-Ib, aph(3’)-Ia, aph(3’’)-Ib, aph(6)-Id, and aac(3)-Ia* **(Figure 6)**. These genes encode the most prevalent aminoglycoside modifying enzymes conferring resistance to tobramycin, amikacin, and kanamycin [52]. These genes were more abundant under laboratory-selection antibiotic concentration (50 mg L^−1^), helping bacteria like *Microbacterium* sp., *Bosea* sp., and *Zoogloea* sp. to resist this condition. *aph(3*″*)-Ib*, and *aph(6)-Id* have been described, complete or in part, within plasmids, integrative conjugative elements, and chromosomal genomic islands [53, 54]. As a consequence of the dissemination of this DNA fragment, the *aph(6)-Id* and *aph(3*″*)-Ib* genes are found in both Gram-positive and Gram-negative organisms [55].

The presence of exDNA and an antibiotic can promote the structural integrity of biofilms [56], hence facilitating bioaggregation and biomass development in the reactor. Microorganisms in the test reactor aggregated easier than in the control reactor **(Figure S13)**. Corno et al. [57] have observed that antibiotics (levofloxacin, tetracycline, and imipenem) can increase the cellular clustering as microcolonies and bioaggregates by 3% (at low antibiotic loading) to 20-25% (at high antibiotic loading) in artificial lake water bacterial communities studied in chemostats. According to Das et al. [58], adding exogenous DNA in pure cultures increases bacterial adhesion and broth viscosity. This was facilitated with an amendment of Ca^2+^ through acid-base interactions and cationic bridges.

Biomass growth under high antibiotic concentrations mainly resulted from the presence of resistant bacteria already in the inoculum rather than a consequence of transformation. The qPCR showed that the plasmid remained in the test chemostat even after 8 days (i.e., after 8 HRTs, 12 number of generations) after stopping its spiking (**Figure 4**). The plasmids remained either transformed inside the predominant bacterial populations selected or adsorbed to EPS. Proximity-ligation Hi-C sequencing was used to quantify the interactions between the pool of genomic and mobile DNA molecules in the same cell within the microbial community, as discussed hereafter.

### 4.2. Microbial community and ecological dynamics of plasmid transfer

It is known from pure cultures of *Streptococcus pneumoniae* and *Legionella pneumophila*, that the exposure to aminoglycoside and fluoroquinolone antibiotics induces their genetic transformability (*i*.*e*., competence state) as a result of genotoxic stress [59–61].

Kanamycin promotes competence by inducing decoding errors during translation into aminoacids. It inhibits protein synthesis by binding to the A site of 16S rRNA in the 30S ribosomal subunit and generates misfolded proteins that activate the serine protease HtrA. This triggers a cascade of reactions involving interactions between competence-stimulating proteins (ComC, ComAB, CSP, and others) that eventually launches the competence state [59, 62].

In this study, microorganisms from different classes **(Figure S7)** displayed plasmid-host interactions. *Alphaproteobacteria, Betaproteobacteria, Bacteroidetes*, and *Actinobacteria* contain one or more species that are naturally competent [7]. Experimental demonstrations of natural competence have so far been limited to only a few dozen of species scattered across the bacteria tree and investigated in pure cultures. Such culture-based assays are only conducted with species where a selectable genetic marker is available and cannot help discover competence in uncharacterized populations present in microbial communities that generally do not grow on agar plates [7]. Our approach combining mixed-culture biotechnology and Hi-C metagenomics sequencing successfully uncovered transformability in a microbial community of activated sludge. Further research on competence genes expression and proteome analysis would elucidate the mechanisms by which bacteria have taken up exDNA.

Most of the identified transformed bacteria were Gram-negative (*Bosea* sp., *Runella* sp., *Gemmobacter* sp., and activated sludge floc former *Zoogloea* sp.) and one Gram-positive (*Microbacterium* sp.). Gram-negative bacteria comprise an outer membrane that protects them against the antibiotic and makes them more challenging to kill. Gram-positive ones have a thick peptidoglycan layer that absorbs antibiotics and detergents easier, leading to faster cell death and slower development of resistance [63]. We recently showed that most microorganisms co-localizing MGEs and ARGs in a full-scale WWTP were Gram-negative [4].

A diverse microbial community like activated sludge is composed of many potential hosts that encompass a diversity of mechanisms to maintain and transfer plasmids, including donor-mediated conjugation once the plasmid is transformed [64]. Conjugative plasmid transfer spreads ARGs even in the absence of antibiotic pressure, as reported from investigations of the gut microbiota [65]. Here, as displayed in **Figure 6**, at high antibiotic pressures, there were more hosts containing aminoglycoside resistance genes located in plasmids. Tracking the temporal dynamics of plasmids uptake via transformation together with the more recurrent microbial hosts and spreaders is an important outcome for evaluating the risks associated with the transformation of exogenous (synthetic) MGEs, on top of conjugative events. With this study, we provide the tools to achieve such target in complex microbial communities

### 4.3. Limitations and alternatives of Hi-C sequencing for detecting transformation events

When building MAGs from microbial communities, microdiversity within a genus or species can challenge the taxonomic affiliation resolution, such as observed for the closely related *Runella* spp. Besides a taxonomic classification problem, binning can also be affected in such situation. When multiple populations of a single genus are present in a microbial community, parts of a contig can derive from one strain while other parts from other strains can coassemble [37].

Detecting specific fractions of the spiked plasmid interacting with genome clusters (or MAGs) of populations from the mixed culture was challenging. First, commercial and publicly available bioinformatics platforms (ProxiMeta) and computational tools (bin3c) were not sensitive enough to track individual transfer events, while very useful for general resistome and plasmidome to host linkage. Second, only 17.9% of the contigs of the metagenome assembly could be sorted into MAGs (**Figure S14-S15)**. This leaves 82.1% of the genetic information unanalyzed, potentially containing information about the plasmid-host. Therefore, manual data curation involving all discordant reads analysis and cluster linkage was done. Discordant read analysis was advantageous to compare all contigs of the metagenome interacting with contigs affiliating with the spiked plasmid sequence, bypassing the binning limitation.

epicPCR (emulsion, paired isolation, and concatenation PCR) is an alternative method to Hi-C sequencing for recovering linked phylogenetic and functional genetic information from millions of cells in a single analysis [66]. epicPCR is limited by the requirement of prior knowledge of sequences of the target genes before performing the qPCR and by the biases introduced by amplifying the 16S rRNA gene [67, 68]. To increase sensitivity, combining Hi-C sequencing and epicPCR would be interesting for identifying transformation events in mixed cultures and identifying new potential natural competent bacterial species.

### 4.4. Natural transformation and rolling-circle plasmids as models

In this transformation experiment of the mixed culture, we used the rolling-circle replication (RCR) plasmid pBAV1K-T5-GFP. This plasmid is a suitable model plasmid for studying natural transformation in microbial communities through its capacity to replicate in both Gram-positive and Gram-negative microorganisms [17]. RCR is one of the simplest mechanisms adopted by some plasmids, which relies on a sequence-specific cleavage. This generates a nick in the double-strand origin of one of the parental DNA strands by an initiator Rep protein (3’-OH end), allowing DNA polymerases to initiate the leading strand replications [69] and circumventing the primer RNA synthesis used in the canonical theta replication.

There are multiple ways by which the spiked plasmid could be incorporated into the genomes of the competent bacteria of the activated sludge. The canonical way is via double-stranded exDNA uptake through the outer membrane, the periplasm, and internalized as single-stranded DNA through the inner membrane (in Gram-negative). In Gram-positive, the exDNA must cross a thick peptidoglycan layer [70]. Methylation plays a protective role in bacteria for avoiding exogenous DNA influx from bacteriophages [71, 72]. The plasmid DNA spiked in the test chemostat was produced in an *E. coli* strain with its dam/dcm methylases activated. However, DNA is usually single-stranded inside the cytoplasm in natural transformation and thus not a target for most restriction enzymes [73].

Double-stranded plasmid DNA could also be transformed via σ^S^ regulation through ABC transporters [74]. In natural transformation in *E. coli*, dsDNA passes across the outer membrane through an unknown pore. This way, dsDNA could transform and not be genome integrated but maintained episomally. This research focused on the possibility of exDNA being integrated into the genome. However, from the microbial ecology point of view, more research should be done on how bacteria exchange genetic information in samples from complex systems to understand microbial evolution.

### 4.5. Outlook

Overall, among the three main mechanisms of HGT, transformation rarely occurs between bacterial species to transfer drug resistance genes [75] when compared to more efficient processes such as conjugation, since conjugative plasmids account for half of all plasmids, and these can be broad host range [64, 76, 77]. Antibiotic concentrations in the environment range typically from 0.01 µg L^−1^ in the sea to 0.1 µg L^−1^ in rivers, 1 µg L^−1^ in treated municipal sewage, 10 µg L^−1^ in untreated municipal sewage, and up to 100-10’000 µg L^−1^ found in untreated hospital effluents and industrially polluted surface water [78]. Here, transformation was detected under high antibiotic concentration (>50 mg L^−1^), *i*.*e*., resembling highly concentrated antibiotic streams or antibiotic administration in the gut and polluted industrial waters. Such antibiotic level was used to detect a clear microbial community response from the experimental noise. The consequences of single transformation events may be vast [78], potentially inducing a severe medical issue by developing the so-called “superbugs” or bacteria resistant to two or more antibiotics. The following research should focus on the environmental conditions (nutrient limitation, temperature shifts, linear vs. plasmid DNA, methylation patterns, among others) that trigger competence, exDNA uptake, and exchange within microbial communities.

## 5. Conclusions

We identified that microorganisms in a mixed culture enriched from activated sludge could take up and get transformed by synthetic plasmids present in their wastewater environment, provided a selection pressure is present, like a high antibiotic concentration. The result of this work involving quantitative mixed-culture biotechnology and Hi-C sequencing led to the following additional main conclusions:

1. The spike of plasmid DNA helped the biomass adapt and to have a higher yield under high antibiotic concentrations (> 50 mg L^−1^).
2. The plasmid DNA accumulated in the test chemostat even when its spiking was stopped for 8 days, by either uptake inside bacterial cells and/or adsorbed to the EPS.
3. High kanamycin concentrations (50-100 mg L^−1^) promoted the mobility of aminoglycoside resistance genes among bacterial hosts.
4. Environmental antibiotic concentrations (≤2.5 mg L^−1^) did not induce detectable transformation events, while lab-selection concentrations (50 mg L^−1^) did.
5. The main hosts containing the spiked synthetic plasmid pBAV1K-T5-GFP in the activated sludge were the Gram-negative bacteria *Bosea* sp. and *Runella* sp. (accompanied by *Gemmobacter* sp. and the well-known activated sludge floc former *Zoogloea* sp), and the Gram-positive *Microbacterium* sp..
6. The outcomes are important for not only the science but also the mitigation of the transfer of antibiotic resistance and foreign genetic elements emitted in wastewater catchment areas, especially in regions without wastewater treatment infrastructure or where industrial wastewater is not treated.

## Data availability

Metagenome sequencing and amplicon sequencing data were deposited in the NCBI database with the BioProject ID: PRJNA868937.

## Authors’ contributions

DCF designed the study with inputs from MvL, TA, and DGW. DCF performed the experimental investigations. DCF wrote the manuscript with all authors’ direct contributions, edits, and critical feedback.

## Acknowledgments

We thank Riko Kuriki for the control strain development and microscopy pictures. We warmly acknowledge Marinka Almering and Zita van der Krogt for helping us build a biosafety level II facility to perform this experiment. This work is part of the research project “Transmission of Antimicrobial Resistance Genes and Engineered DNA from Transgenic Biosystems in Nature” (TARGETBIO) funded by the programme Biotechnology & Safety Program of the Ministry of Infrastructure and Water Management (grant no. 15812) of the Applied and Engineering Sciences (TTW) Division of the Dutch Research Council (NWO).

