## Supplementary Material for "Catch me if you can: Capturing extracellular DNA transformation in mixed cultures via Hi-C sequencing"

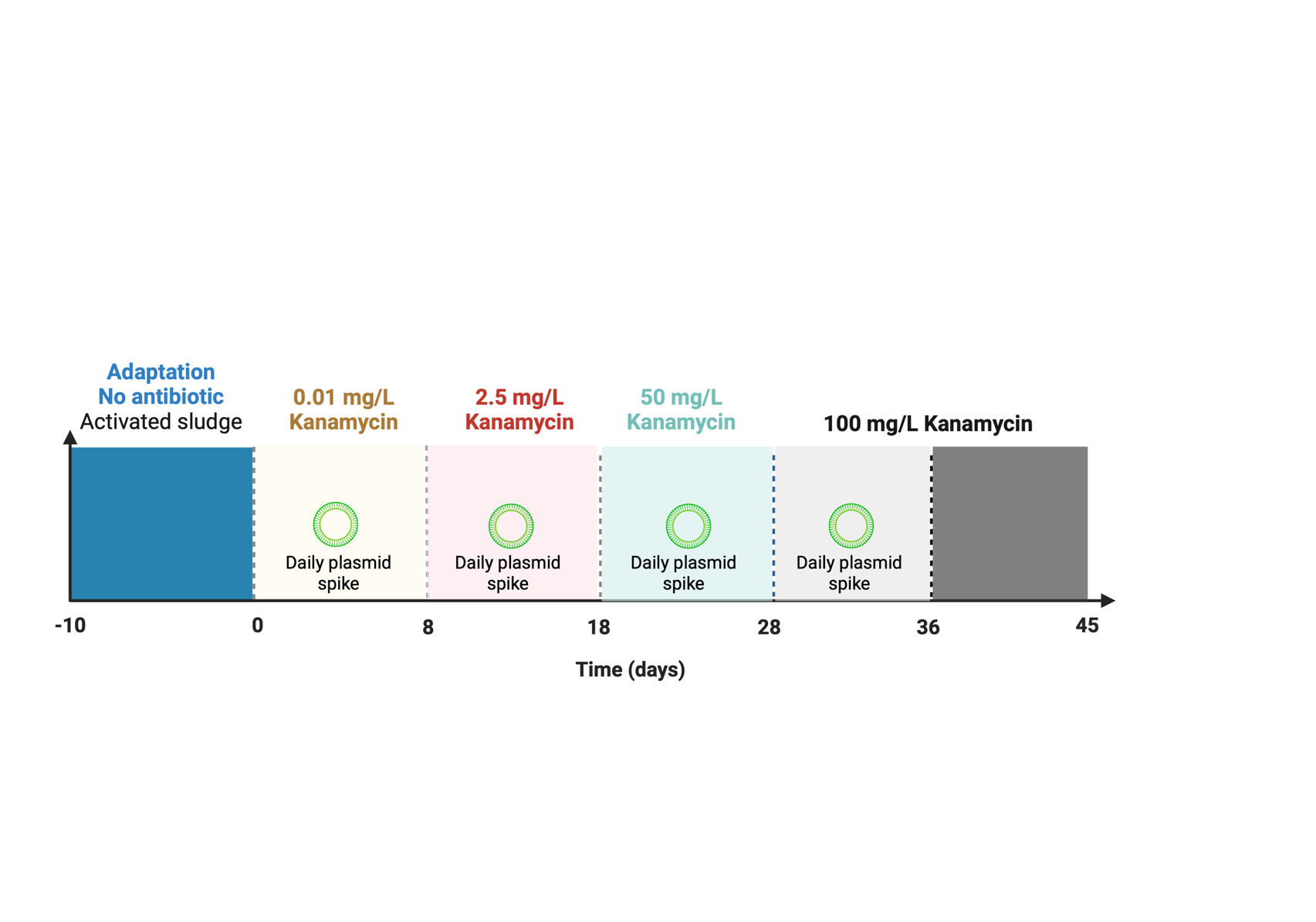

**Figure S1.** Schematic representation of the operation over time in the test reactor. The control reactor was operated equally but without plasmid addition.

**Table S1.** Recipe of influent complex synthetic wastewater receiving reactors test and control. The recipes provide C, N and P for the wastewater preparation. The recipe was prepared in 20-fold concentration to provide total COD and TN concentrations of 600 mg COD L^-1^ and 52 mg TN L^-1^, respectively.

| Component | Complex synthetic WW  Concentration [g L^-1^] |
| --- | --- |
| NaAcetate*3H_2_O | 4.3 |
| NaPropionate | 1.6 |
| (NH_4_)Cl | 1.1 |
| CaCl_2_* H_2_O | 0.35 |
| MgSO_4_ | 0.33 |
| KCl | 0.66 |
| Glucose | 1.9 |
| Starch | 1.4 |
| Peptone | 1.6 |
| Alanine | 0.27 |
| Arginine | 0.26 |
| Aspartic Acid | 0.40 |
| Glutamic Acid | 0.29 |
| Glycine | 0.45 |
| Leucine | 0.16 |
| Proline | 0.19 |
| K_2_HPO_4_ | 0.23 |
| KH_2_PO_4_ | 0.23 |

**Table S2.** Composition of trace element solution, after preparation pH is adapted to 6 using KOH (30% v/v)

| Component | Formula | Concentration [g L^-1^] |
| --- | --- | --- |
| EDTA disodium salt dihydrate | C_10_H_14_N_2_Na_2_O_8_ * 2H_2_O | 16.22 |
| Zinc II sulfate | ZnSO_4_ * 7H_2_O | 0.44 |
| Manganese II Chloride | MnCl_2_ * 6H_2_O | 1.01 |
| Ammonium Iron II | (NH4)2Fe(SO4)2* 6H2O | 7.05 |
| Ammonium Molybdate | (NH4)6Mo7O24* 4H2O | 0.33 |
| Copper II Sulfate | CuSO4* 5H2O | 0.31 |
| Cobalt II Chloride | CoCl2* 6H2O | 0.32 |

| Gene | Sequence |
| --- | --- |
| *16S rRNA* | ACTCCTACGGGAGGCAGCAGTGGGGAATATTGCACAATGGGCGCAAGCCTGATGCAGCCATGCCGCGTGTATGAAGAAGGCCTTCGGGTTGTAAAGTACTTTCAGCGGGGAGGAAGGGAGTAAAGTTAATACCTTTGCTCATTGACGTTACCCGCAGAAGAAGCACCGGCTAACTCCGTGCCAGCAGCCGCGGTAAT |
| *GFP* | TTCAATGCTTTTCCCGTTATCCGGATCATATGAAACGGTATGACTTTTTCAAGAGTGCCATGCCCGAAGGTTATGTACAGGAACGCACTATATCTTTCAAAGATGACGGGAACTACAAGACG |
| *Kan^R^*  *(aph(3’)-IIIa)* | CACTTACTTTGCCATCTTTCACAAAGATGTTGCTGTCTCCCAGGTCGCCGTGGGAAAAGACAAGTTCCTCTTCGGGCTTTTCCGTCTTTAAAAAATCATACAGCTCGCGCGGATCTTTAAATGGAGTGTCTTCTTCCCAGTTTTCGCAATCCACATCGGCCAGATCGTTATTCAGTAAGTAATCCAATTCGGCTAAGCG |

**Table S3.** 16S rRNA, GFP and Kan^R^ synthetic DNA fragments used to generate standard curves for qPCR

**Pure culture laboratory transformation**

**Transformation of *E. coli K12* by electroporation**

Electroporation was performed by preparing electrocompetent *E. coli* K12 cells by inoculating *E. coli* K12 in LB medium overnight at 37˚C 180 rpm. 100 mL of fresh LB medium was prepared in 500 mL shake flask and the culture grown overnight was added to a final OD_600_ of 0.02 and further incubated at 37˚C 200 rpm until it reached OD_600_ 0.5-1.0. Cells were collected at 2000 x g at 4˚C for 15 min. Cells were washed three times using 10% of the original culture volume with precooled filter sterile 10% glycerol. 50 µL of washed cells and 1 ng of pBAV1K-T5-gfp were transferred to prechilled Gene Pulser®/Micropulser™ electroporation cuvettes (Bio-Rad). After 2 min of incubation on ice, an electrical pulse of 12.5 kV·cm^-1^ was added, resulting in a time constant of 4.3 to 4.5 ms. Immediately after the pulse delivery, 1 ml of prewarmed LB medium was added to cells. After incubation at 37˚C for 45 min, the cells were plated on LB agar plates containing 50 µg mL^-1^ kanamycin for selection of transformed cells containing pBAV1K-T5-gfp overnight at 37˚C.

**Transformation of *B. subtilis 168* by starvation-induced competence**

Transformation of *B. subtilis* str 168 was carried out following a published protocol [18]. Cultures of *B. subtilis* 168 were grown overnight at 37˚C 225 rpm. In a freshly prepared pre-warmed 15 ml SM1, 1 mL of the overnight culture was transferred and diluted to a final OD_600_ of 0.4-0.6 and incubated at 37˚C 225 rpm for 5 hours. Once the culture reached the stationary phase (OD_600_ of 2.0-2.8), equal volume of pre-warmed SM2 was added and incubated for 90 min at 37˚C for 2h. 500 µL of cell culture was combined with 5 µl (100-500 ng) plasmid DNA and incubated for 30 min at 37˚C at 180 rpm. 300 µL of LB was added and further incubated for 30 min at 37˚C 180 rpm. 100 µL of cells were plated onto LB agar plates containing 50 µg mL^-1^ of kanamycin and grown overnight at 37˚C.

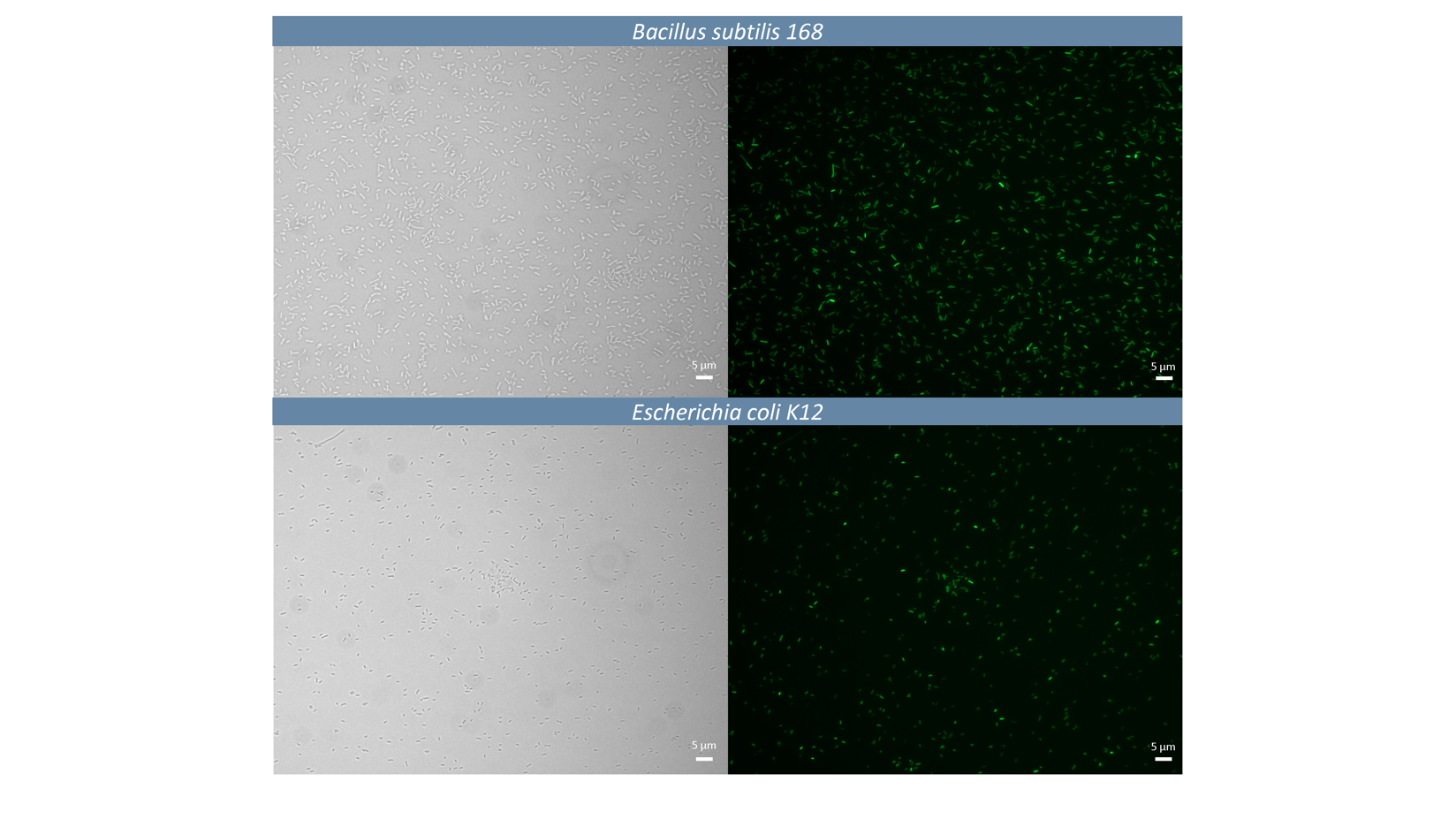

**Figure S2.** Microscopic pictures of gram-positive *Bacillus subtilis 168* **(upper panel)** and gram-negative *Escherichia coli K12* **(bottom panel)** under bright field (**left figures)** and under fluorescent microscopy **(right figures)** as a consequence of GFP expression

**Chemostat operation parameters**

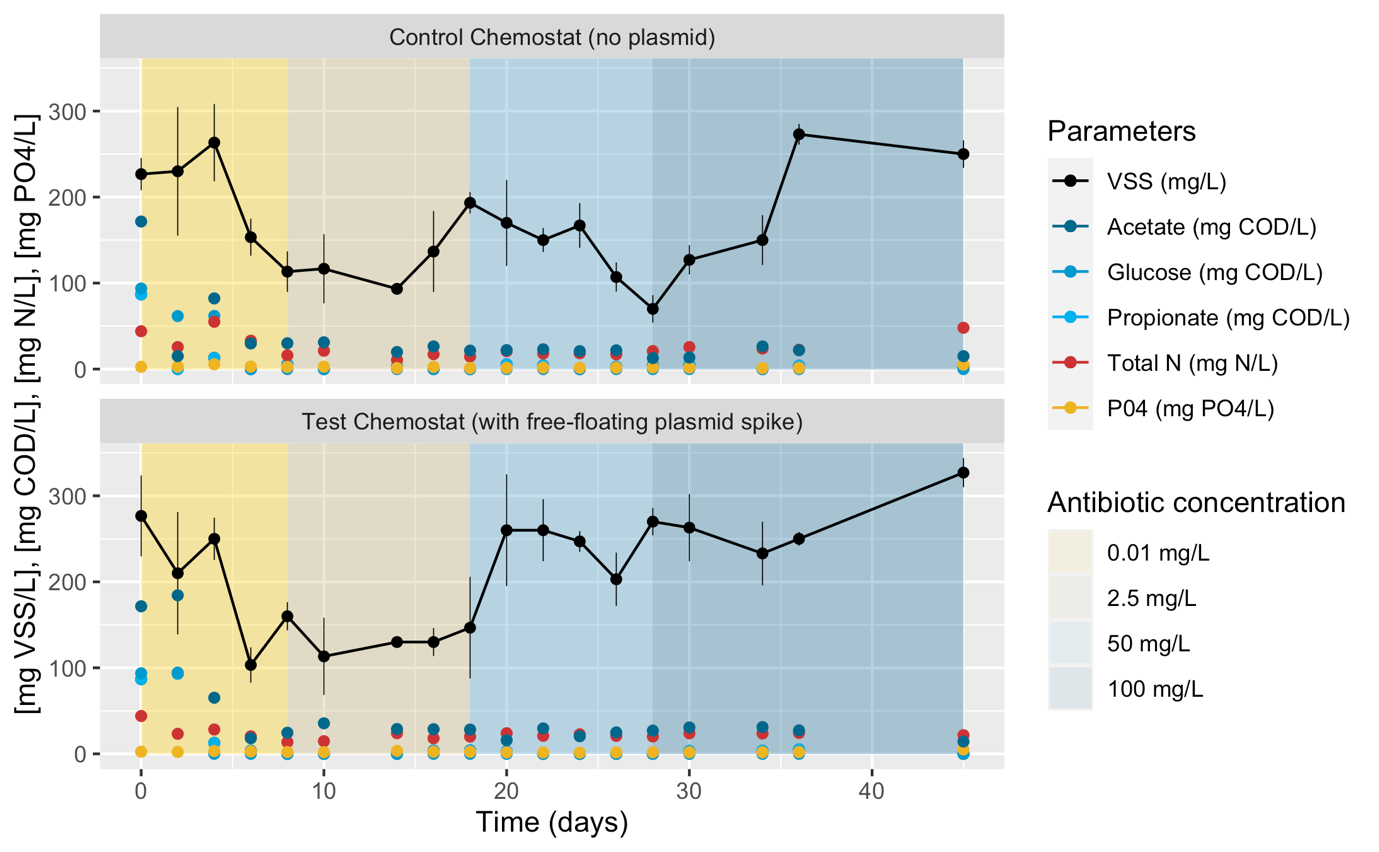

**Figure S3.** Daily-averaged values for volatile suspended solids (VSS), volatile fatty acids (VFAs), glucose, total nitrogen and phosphate from both reactor control and reactor with free-floating plasmid over the whole operation time the experiment was conducted (45 days). Kanamycin concentrations are displayed as background-colored sections: 0.01 mg L^-1^, 2.5 mg L^-1^, 50 mg L^-1^, 100 mg L^-1^.

**Aminoglycoside resistance genes alignment and tree**

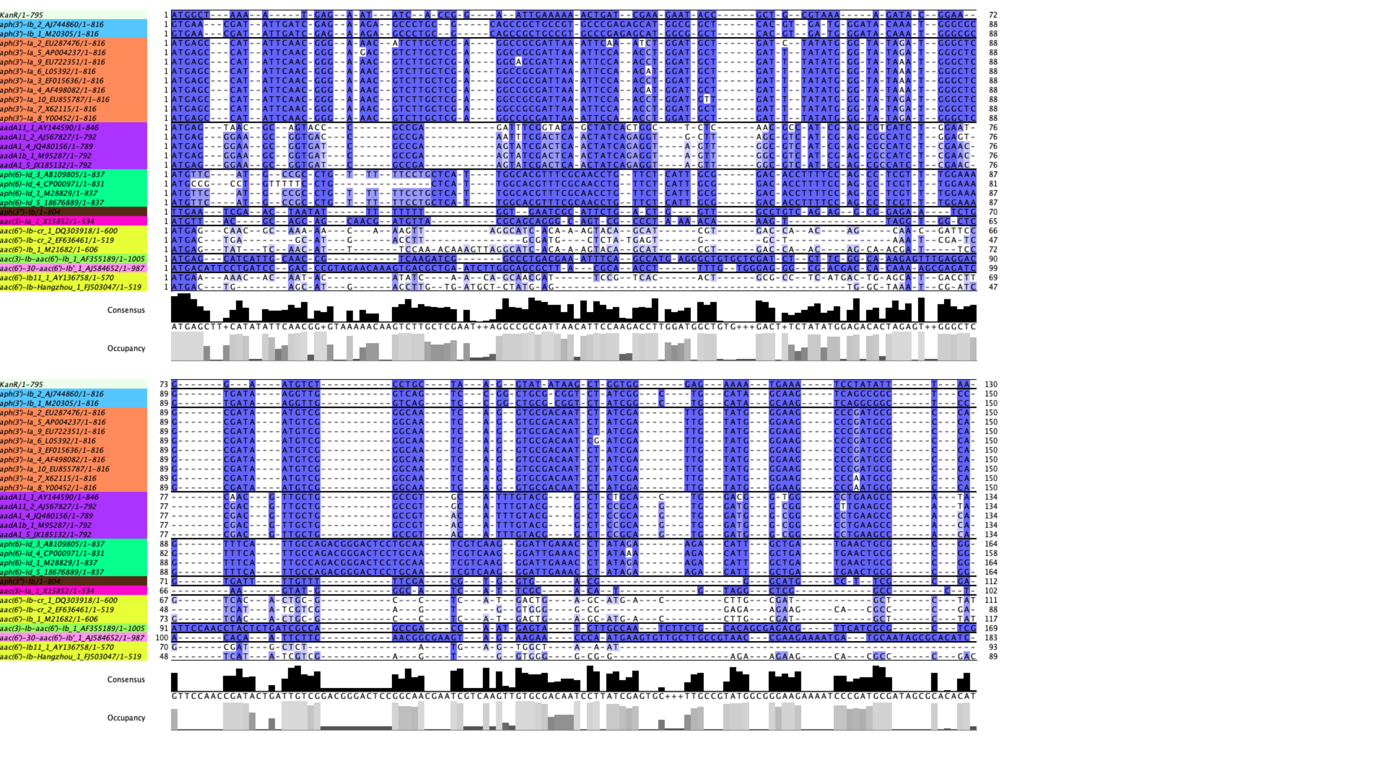

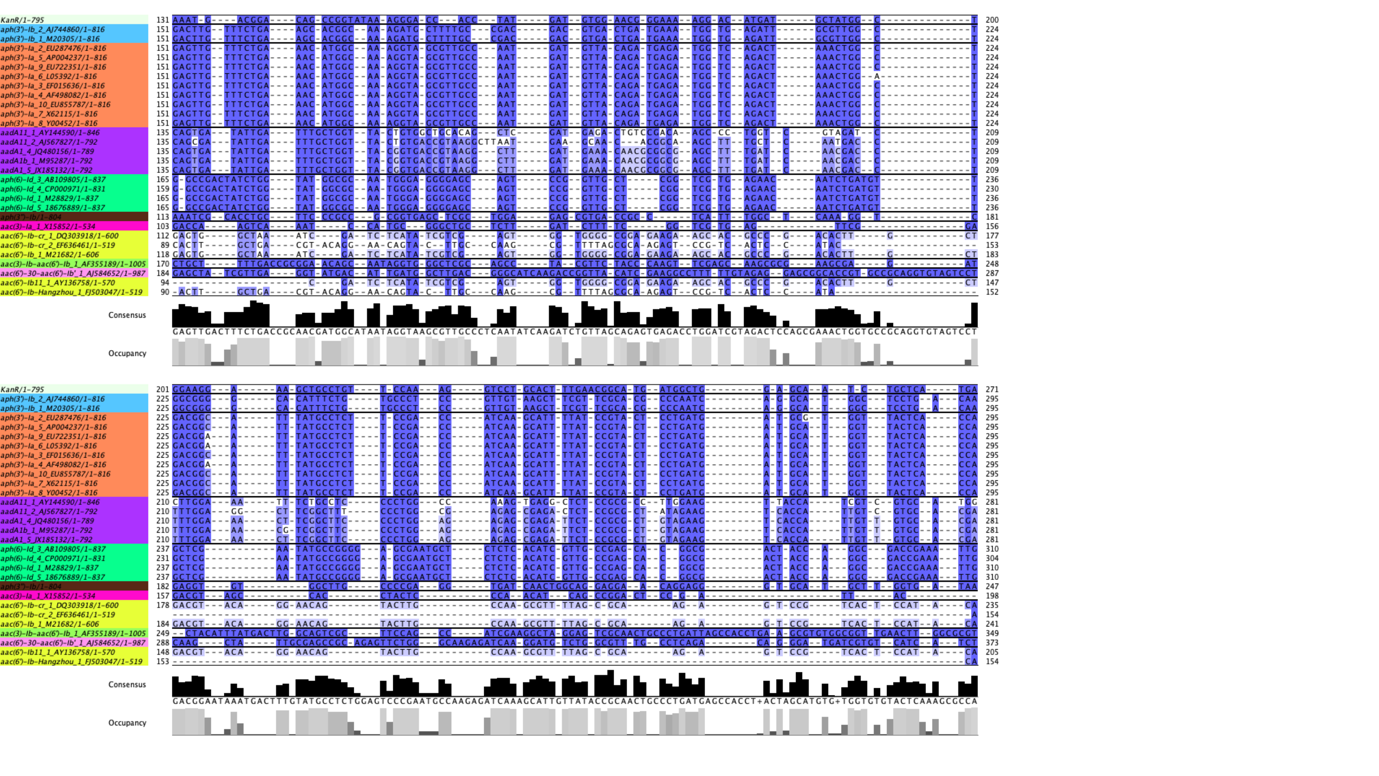

**Figure S4.** Alignment visualization (Jalview-tcoffee) from the aminoglycoside resistance genes (and variants) found inside bacterial hosts in **Figure 6.**

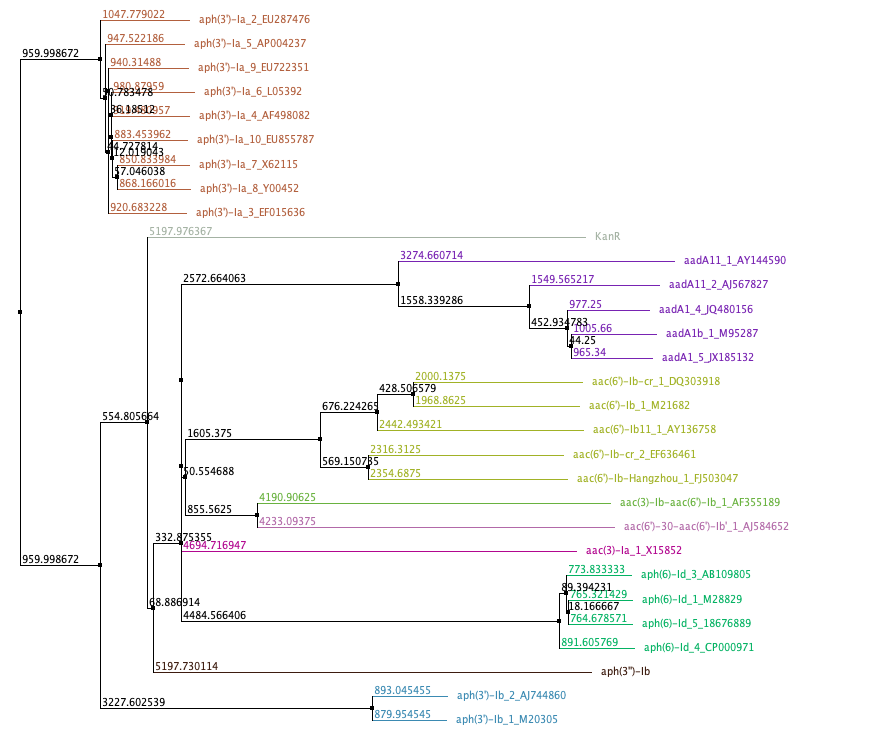
**Figure S5.** Neighbor-joining tree, calculated from the input alignment, is being used to cluster sequences in the main alignment window. The input alignment consisted of the aminoglycoside resistance genes (and variants) found inside bacterial hosts in **Figure 6.**

**Discordant read analysis of individual events**

**Discordant read analysis workflow**

The first thing to identify discordantly mapped reads was to determine where the plasmid contigs were by BLASTn (e-value 1^-20^, coverage 90%) using the genes from the pBAV1K-T5-GFP as query and the assembly as database. From the results, a list containing all the plasmid contigs is created that will be used afterwards.

Then, we need to obtain a file from the sam file from aligning the Hi-C reads against the assembly, where paired-end reads are aligned in different contigs. For this end, awk was used to generate a bam file by extracting mate reads mapped on different contigs using the expression: awk '($3!=$7 && $7!="=")'.

awk '($3!=$7 && $7!="=")' hic_assembly.sam > hic_disc_map.sam

samtools view -S -h -b hic_disc_map.sam > hic_disc.bam

Then, we sort the result file by name:

samtools sort -n hic_disc.bam > hic_disc_sort.bam

Then, we select the entries only containing discordant reads where the contigs containing the information of the plasmid are involved:

samtools view hic_disc_sort.bam |fgrep -w -f list_of_plasmid_contigs.txt > hic_disc_plasmid_contigs.txt

Where the **list_of_plasmid_contigs.txt** are the contigs coming as a result of the BLASTn contigs vs. plasmid nt database:

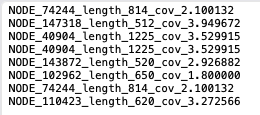

To extract the sequences from, for example the contigs 40904_length_1225 (corresponding to GFP):

grep "NODE_40904_length_1225_cov_3.529915" hic_disc_plasmid_contigs.txt| awk '{ print $3,$7,$10 }'

This will give us the results in three columns of which 2 nodes interact and the sequence. For example, to extract the reads where contigs aligned in GFP from the alignment:

grep "NODE_40904_length_1225_cov_3.529915\|NODE_147318_length_512_cov_3.949672" hic_disc_plasmid_contigs.bam| awk '{ print $3,$7,$10 }' > GFP_contigs_disc.txt

Looking like:

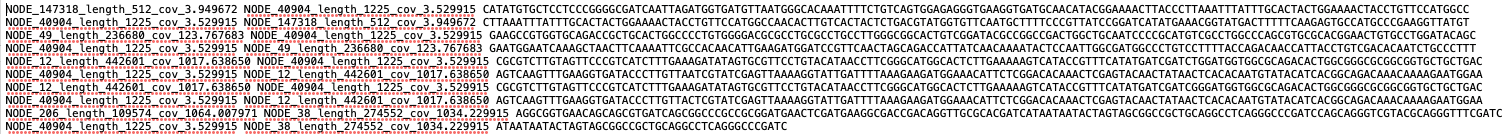

From this file, the NODES (or contigs) containing information from the other parts of the plasmid (not KanR nor GFP genes) were removed in order to avoid quantification of other plasmids that may share genetic information (repA, rrnBT1 terminator and lambda_t0_terminator). Now, a list of contigs containing GFP interacting with multiple other contigs is generated. Then, duplicate contigs were removed in a different file and used to quantify how many times a specific event happened, from which a frequency table could be retrieved and plotted:

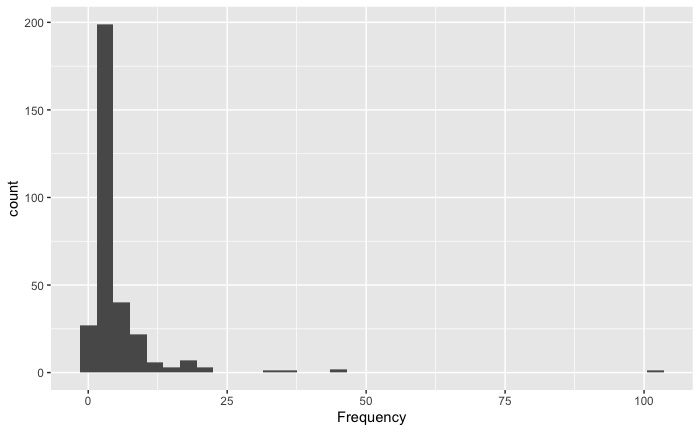

**Figure S6.** Frequency graph showing the number of interactions between contigs from microbial clusters and contigs containing the GFP gene information

We see there are plenty of contigs whose interactions with our reads are higher than 3 (the first column). Those contigs with high frequency (more than 40 interactions) were selected for the next steps.

To further verify that what we have found is correct, a visual analysis of the alignment regions was done. It could be that there are artefacts due to the PCR amplification of the libraries (Hi-C library) so in order to verify if the interactions observed come from real cross-link interaction plasmid-chromosome or if they come due to the PCR artifact, we will create concatenated files and re-align the Hi-C reads against these concatenated files prior visualization. Synthetic constructs consisting of the plasmid sequence, non-coding spacer DNA (1500 bp), and the contigs with higher Hi-C links with plasmid contigs were generated **(Figure S4).**

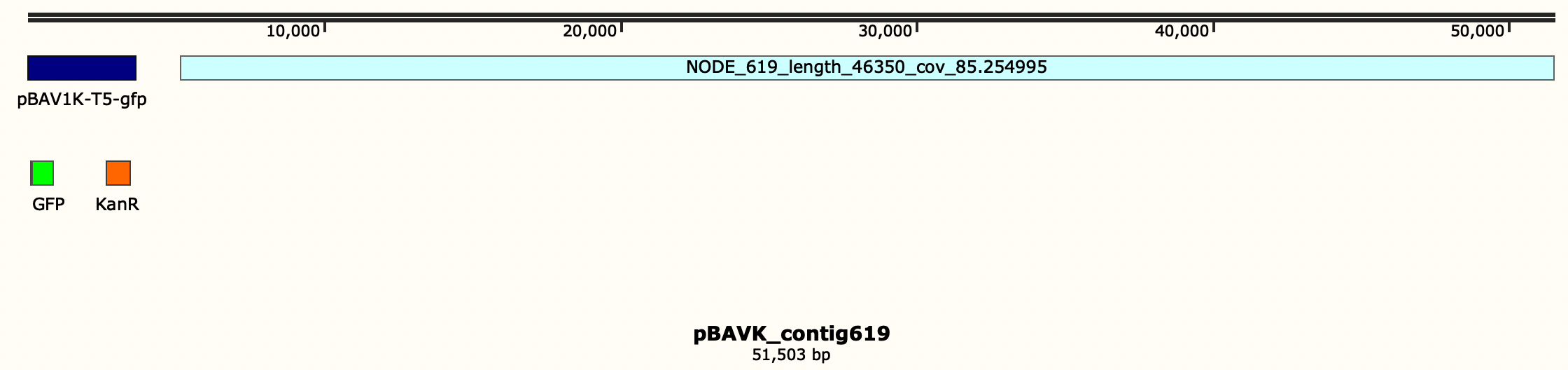

**Figure S7.** Synthetic construct generated containing the sequence of the spiked pBAV1K-T5-GFP plasmid, spacer DNA and the NODE_619 for further visualizing the interactions

Synthetic constructs were aligned with Hi-C forward and reverse reads by bwa-mem v0.7.17-r1188 [25] generating a SAM file, that was converted into a BAM file and then sorted (-n) and indexed with samtools view v1.13 [28]. The resulted file was visualized in IGV [72].

**Taxonomic Tree**

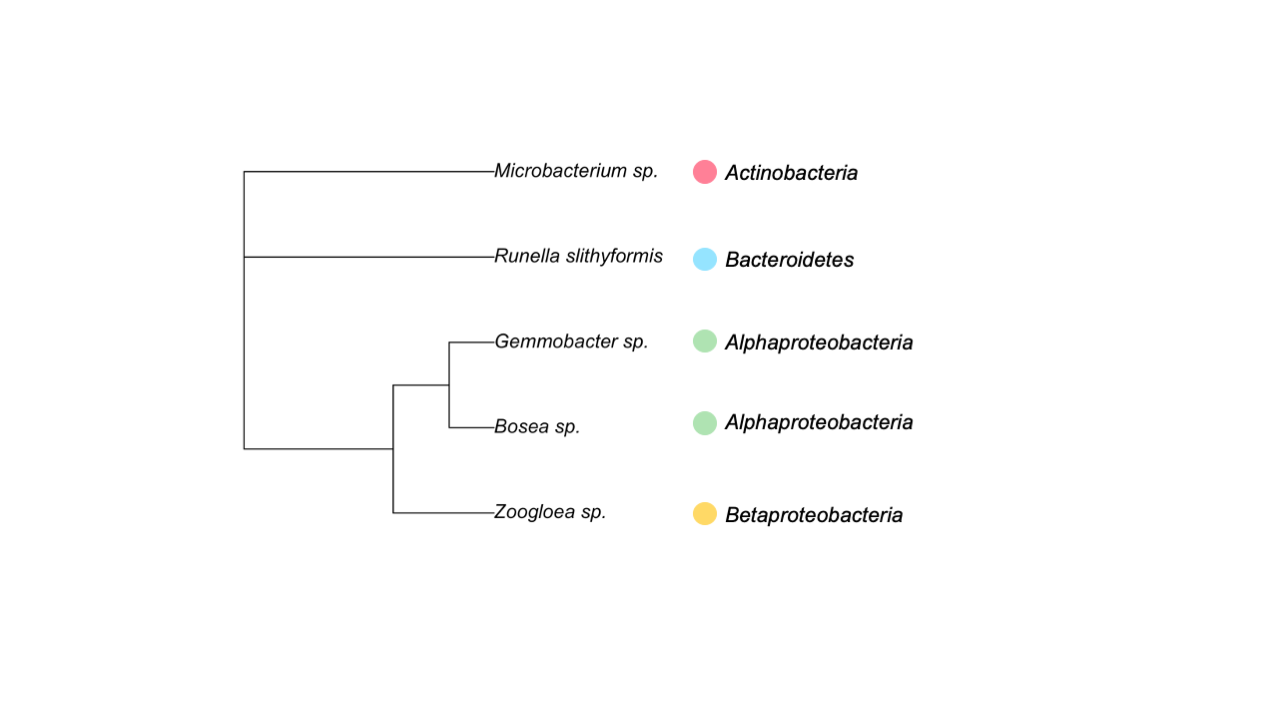

**Figure S8.** Phylogenetic tree of the microorganisms that displayed Hi-C links between their genome clusters and plasmid contigs from the analysis of the discordant reads.

**Discordant reads analysis visualization in IGV**

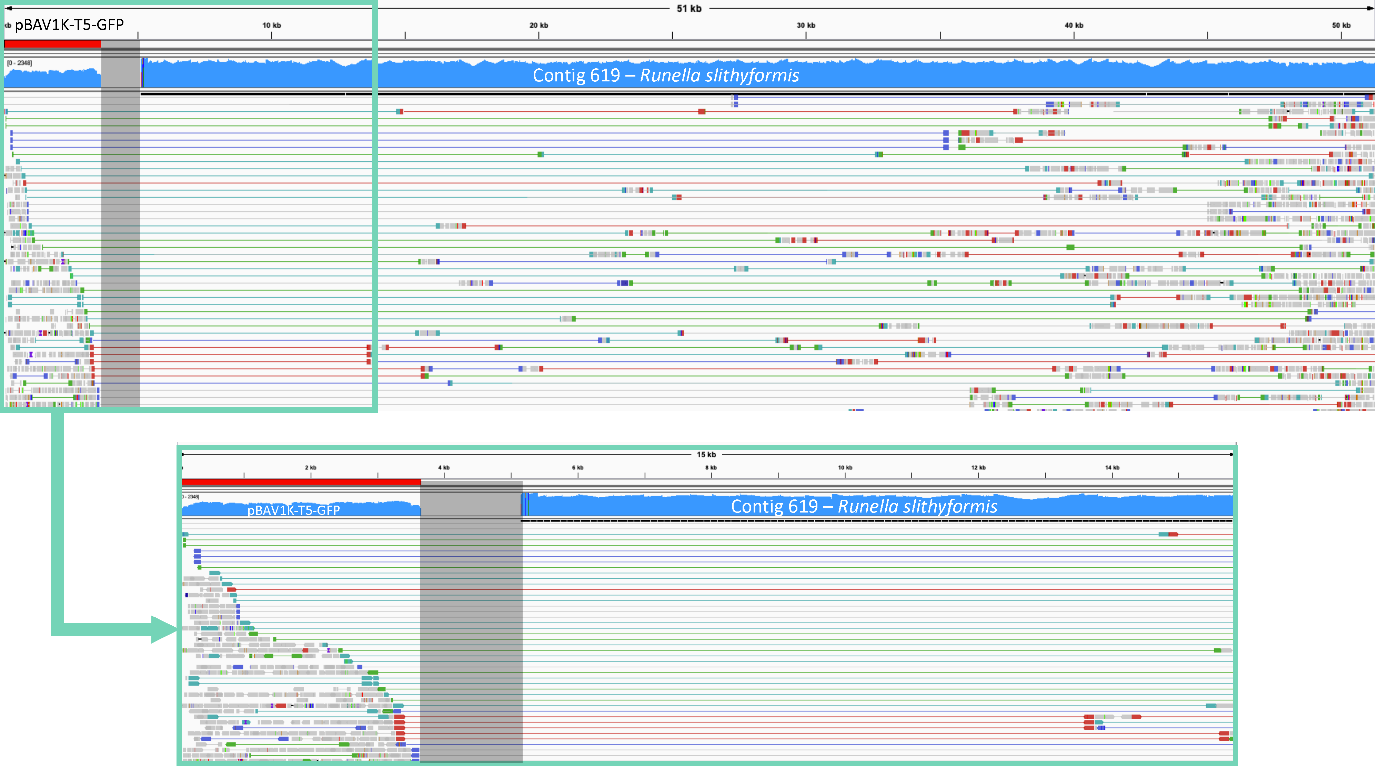

**Figure S9.** Discordant read analysis of contig 619 belonging to the *Runella slythiformis* cluster with the plasmid pBAV1K-T5-GFP.

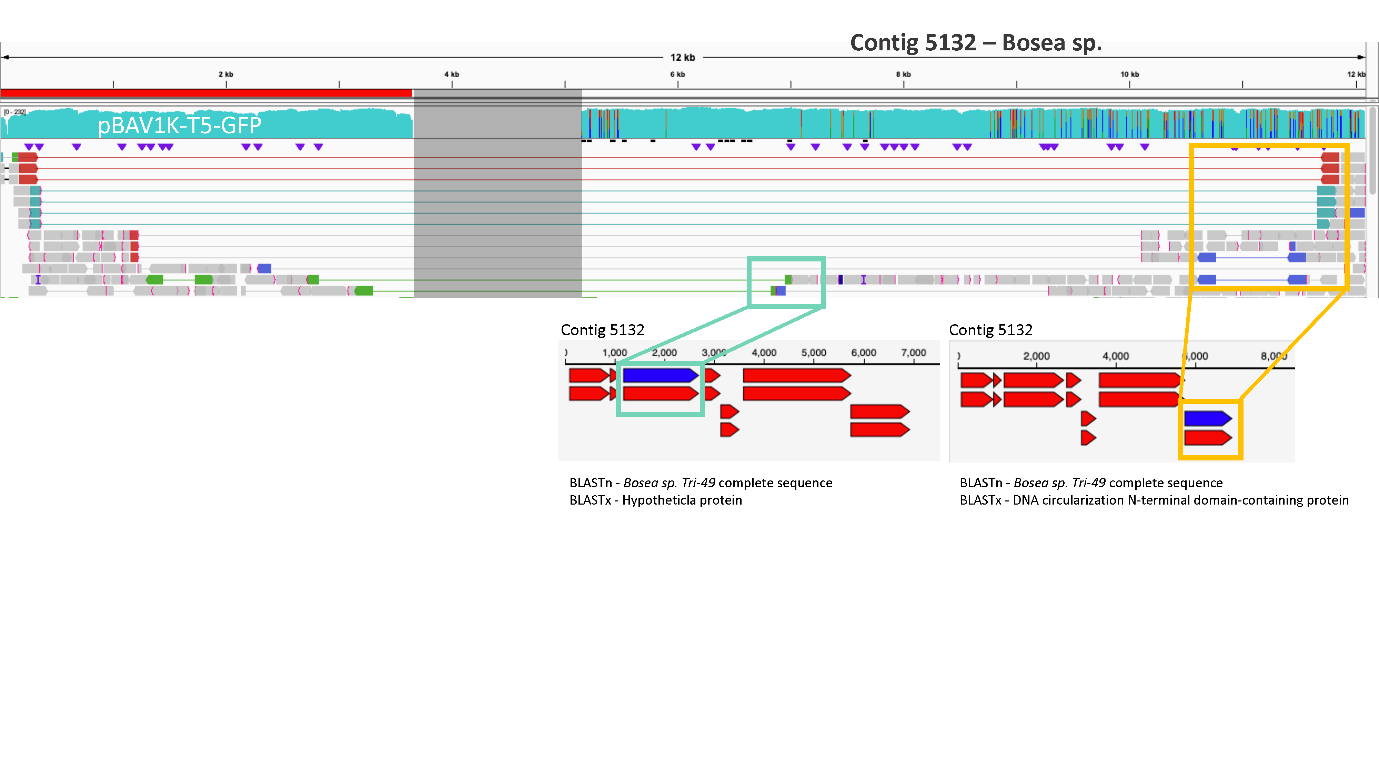

**Figure S10.** Discordant read analysis of contig 5132 belonging to the *Bosea sp.* cluster with the plasmid pBAV1K-T5-GFP. Information of the coding genes where the interaction happened is provided.

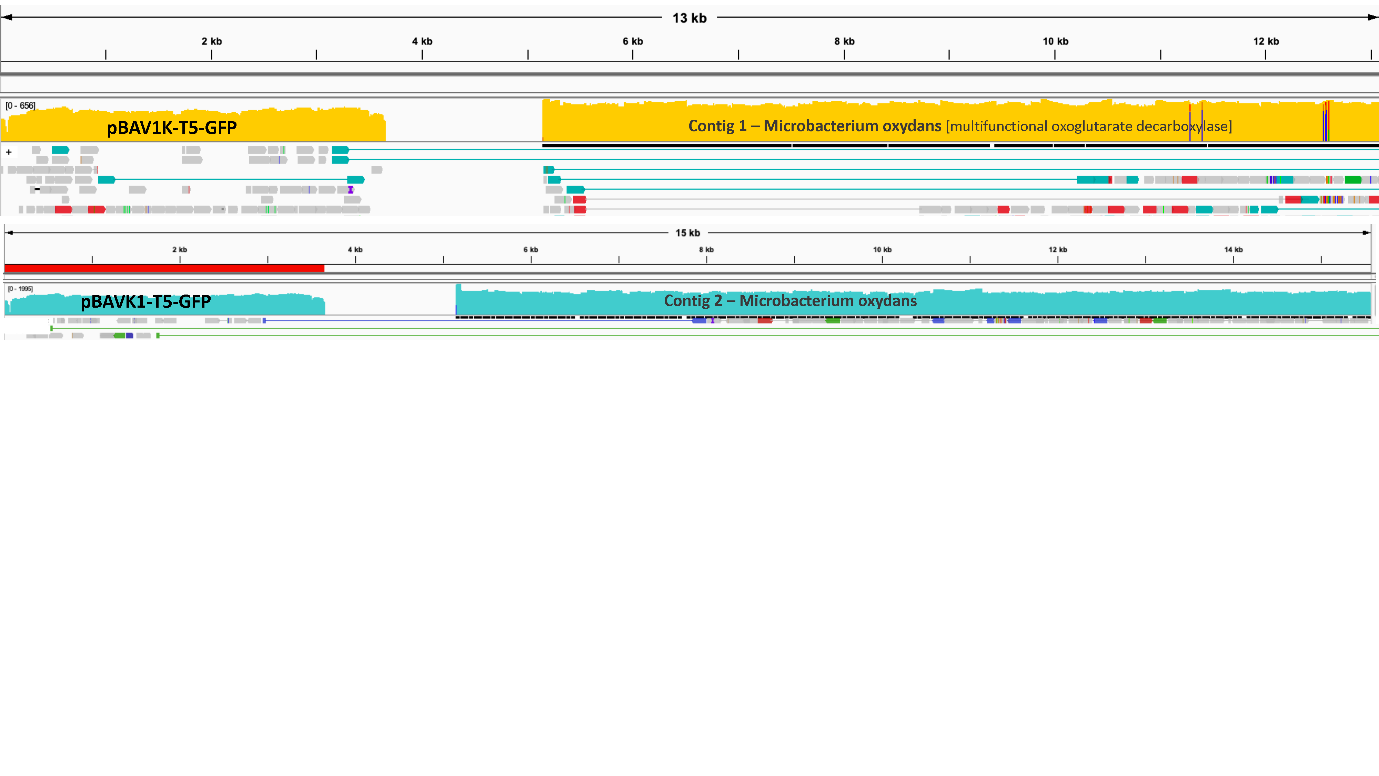

**Figure S11.** Discordant read analysis of contig 1 and 2 belonging to the *Microbacterium oxydans* cluster with the plasmid pBAV1K-T5-GFP. Information of the coding genes where the interaction happened is provided.

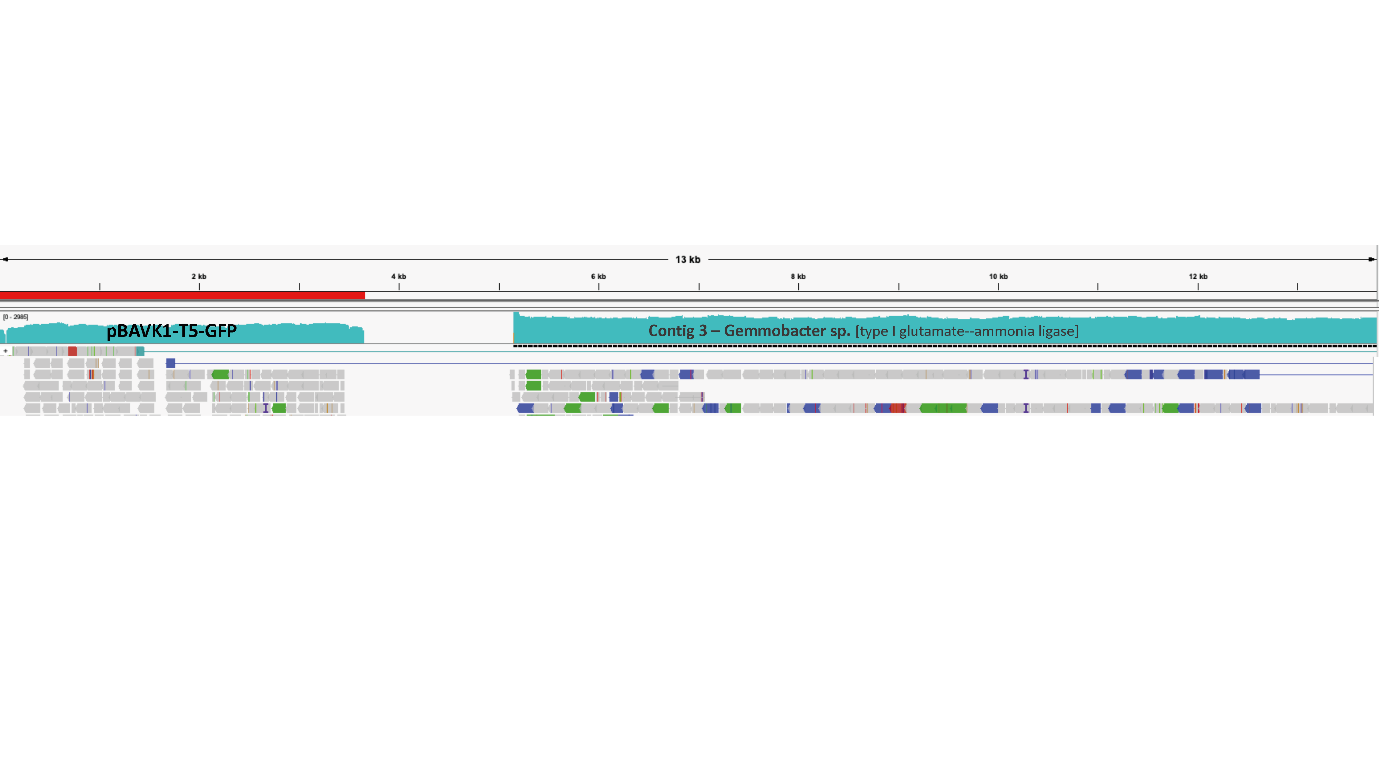

**Figure S12.** Discordant read analysis of contig 1 and 2 belonging to the *Gemmobacter sp.* cluster with the plasmid pBAV1K-T5-GFP. Information of the coding genes where the interaction happened is provided.

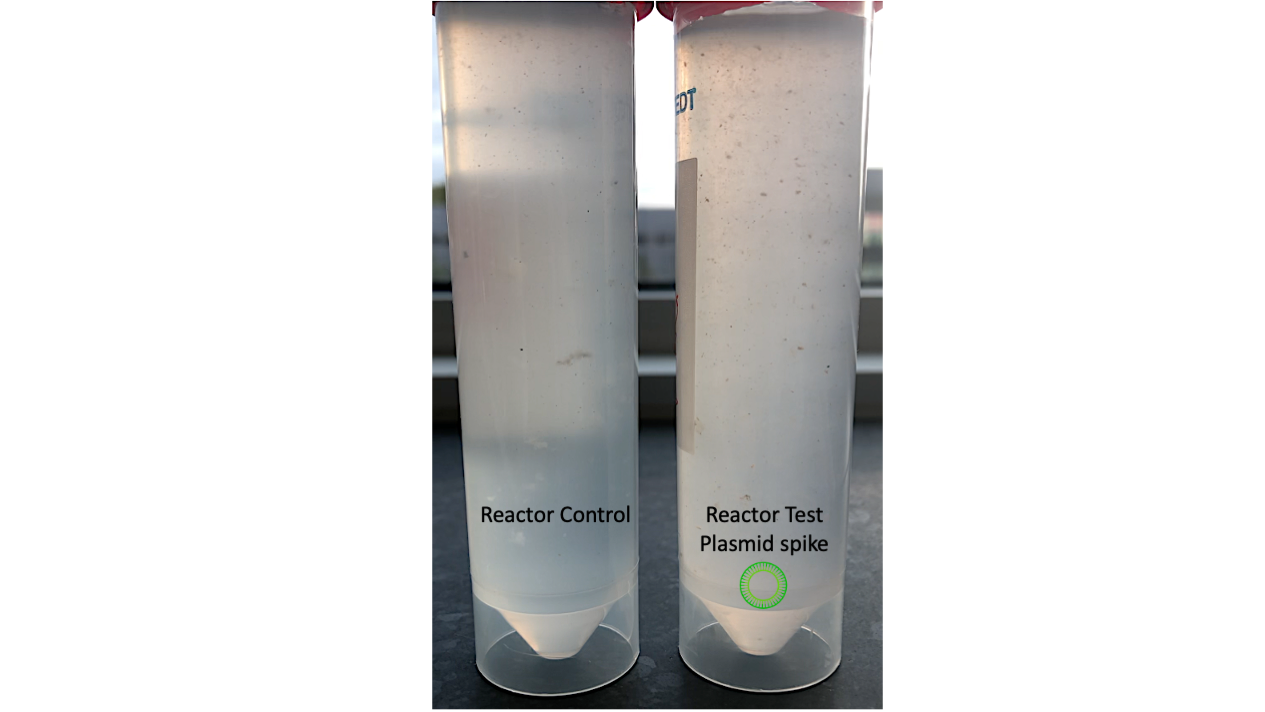

**Figure S13.** Macroscopic morphological visualization of the microbial communities growing in the chemostats. Left tube: Reactor Control (no plasmid spike). Right tube: Reactor test (with free-floating extracellular plasmid addition).

**MAGs recovered**

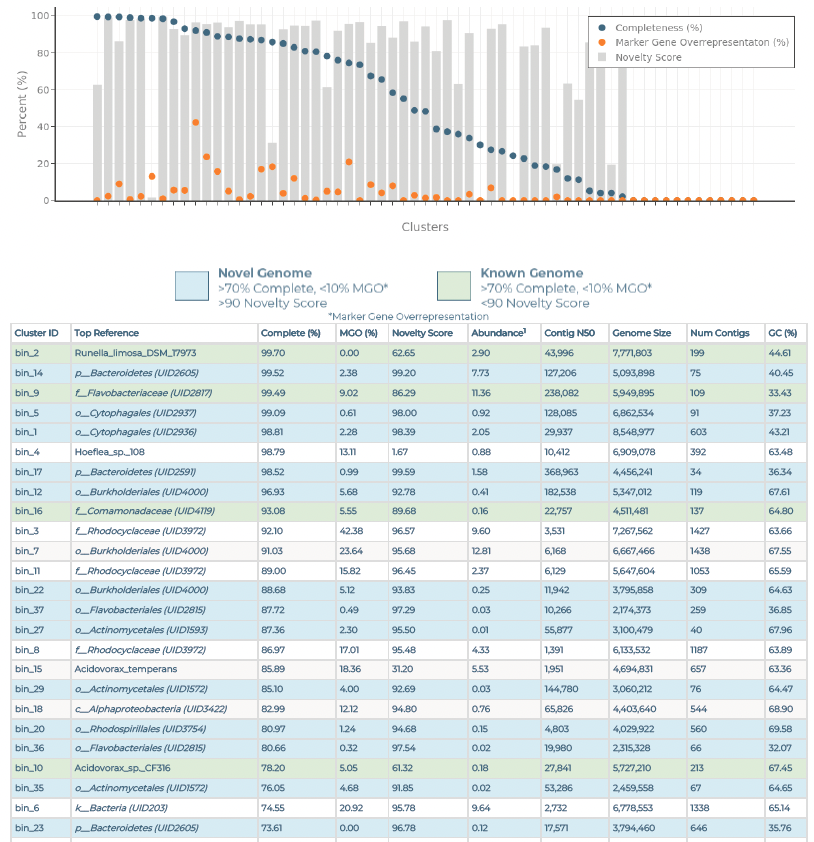

**Figure S14.** Microbial genomes (bins) recovered from the reactor test, day 18 (2.5 mg Kan L^-1^) sorted by highest completeness and lowest contamination.

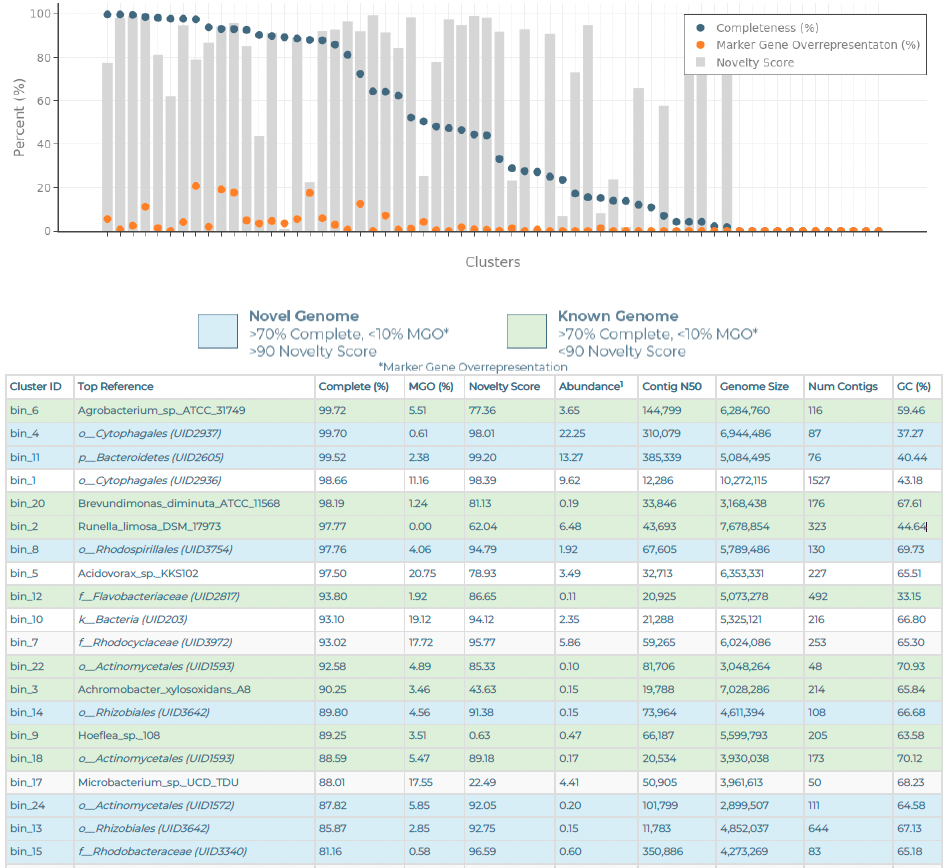

**Figure S15.** Microbial genomes (bins) recovered from the reactor test, day 28 (50 mg Kan L^-1^) sorted by highest completeness and lowest contamination.
